# Modeling the Orthosteric Binding Site of the G Protein-Coupled Odorant Receptor OR5K1

**DOI:** 10.1101/2022.06.01.494157

**Authors:** Alessandro Nicoli, Franziska Haag, Patrick Marcinek, Ruiming He, Johanna Kreißl, Jörg Stein, Alessandro Marchetto, Andreas Dunkel, Thomas Hofmann, Dietmar Krautwurst, Antonella Di Pizio

**Author notes:** These authors contributed equally (Alessandro Nicoli, Franziska Haag). Corresponding Authors, Antonella Di Pizio – Molecular Modeling Group, Leibniz Institute for Food Systems Biology at the Technical University of Munich, 85354 Freising, Germany;, Dietmar Krautwurst - Taste and Odor Systems Reception Group, Leibniz Institute for Food Systems Biology at the Technical University of Munich, 85354 Freising, Germany.

## Abstract

With approximately 400 encoding genes in humans, odorant receptors (ORs) are the largest subfamily of class A G protein-coupled receptors (GPCRs). Despite its high relevance and representation, the odorant-GPCRome is structurally poorly characterized: no experimental structures are available, and the low sequence identity of ORs to experimentally solved GPCRs is a significant challenge for their modeling. Moreover, the receptive range of most ORs is unknown. The odorant receptor OR5K1 was recently and comprehensively characterized in terms of cognate agonists. Here we report two additional agonists and functional data of the most potent compound on two mutants, L104^3.32^ and L255^6.51^. Experimental data was used to guide the investigation of the binding modes of OR5K1 ligands into the orthosteric binding site using structural information from AI-driven modeling, as recently released in the AlphaFold Protein Structure Database, and from homology modeling. Induced-fit docking simulations were used to sample the binding site conformational space for ensemble docking. Mutagenesis data guided side chain residue sampling and model selection. We obtained models that could better rationalize the different activity of active (agonist) versus inactive molecules with respect to starting models, and also capture differences in activity related to minor structural differences. Therefore, we provide a model refinement protocol that can be applied to model the orthosteric binding site of ORs as well as that of GPCRs with low sequence identity to available templates.

## INTRODUCTION

G protein-coupled receptors (GPCRs) are the largest family of membrane proteins in the human genome. Through interaction with their modulators, GPCRs mediate the communication between the cell and the extracellular environment and are, therefore, involved in almost all physiological functions.^1–4^ Commonly, GPCRs are grouped into six classes based on the phylogenetic analysis: A (rhodopsin-like), B (secretin-like), C (metabotropic glutamate receptors), D (pheromone receptors), E (cAMP receptors), and F (frizzled/smoothened receptors).^5–6^ Class A GPCRs consist of over 80% of all GPCRs and are the targets of 34% of all drugs in the market.^7–8^

Class A GPCRs share a basic architecture consisting of a bundle of seven transmembrane α-helices (TM1-TM7) connected by three intracellular loops (ICLs) and three extracellular loops (ECLs), a relatively short N-terminus in the extracellular region, and a short helix 8 connected to the C-terminus in the intracellular module. The ligand-binding domain of class A GPCRs, commonly referred to as the orthosteric binding site, is located in the EC part of the 7TM bundle (made up of residues belonging to TM3, TM5, TM6, and TM7) and has high structural diversity among different receptor subtypes. The 7TM bundle is the most structurally conserved component of the class A GPCR structures, presenting characteristic hydrophobic patterns and functionally important signature motifs.^9–10^

Odorant receptors (ORs), with approximately 400 encoding genes in humans, are the largest subfamily of class A GPCRs.^11–15^ Mammalian odorant receptors are split into two phylogenetically distinct groups, class I and class II ORs, which can be distinguished by some characteristic features that are highly conserved within their sequences.^16–19^ ORs present most of the class A GPCR signature motifs, despite an overall low sequence identity with the non-sensory class A GPCRs.^20–21^ The orthosteric binding site of ORs was also found to coincide with that of non-sensory class A GPCRs.^20–25^

The olfactory system uses a combinatorial code of ORs to represent thousands of odorants: a specific OR type may recognize more than one odorant, and each odorant may be recognized by more than one OR.^26–31^ Despite current efforts in assigning ORs to odorant molecules, or vice versa, in defining the chemical ligand space of individual ORs, only the molecular recognition ranges of a few ORs have been investigated.^27, 32–38^

Structure-based virtual screening campaigns have been successfully applied for GPCR ligand discovery and are always more in use with the recent extraordinary advances in GPCR structural biology.^39^ Currently, no experimental structures of human ORs are available, and homology modeling techniques have been used to rationalize the binding modes of odorant compounds into ORs and discover new OR ligands.^37, 40–44^ AI-based methods are emerging as compelling tools to predict the 3D structure of proteins.^45–47^ During the CASP (Critical Assessment of Structure Prediction) 14 competition, AlphaFold 2 (AF2) was shown to predict the structure of protein domains at an accuracy matching experimental methods.^48^ A database (AlphaFold DB) of over 200 million protein models was released (https://alphafold.ebi.ac.uk/),^49–50^ which expands the coverage for GPCR structures, including odorant receptors.^46^

In this paper, we used both AlphaFold 2 and template-based modeling methodologies for OR5K1 structural prediction. OR5K1 is located on chromosome 3 (3q11.2). It belongs to about 6% of the most abundant human ORs.^51^ OR5K1 has recently been characterized as the specialized OR for the detection of pyrazine-based key food odorants and semiochemicals.^52^ Beyond the olfactory function, physiological functions linked to the extra-nasal expression of OR5K1 cannot be excluded. Indeed, recently it was shown that Olfr177, the mouse ortholog of human OR5K2, which in turn is a homolog to OR5K1, is expressed in the liver and recognizes both pyrazines 2-ethyl-3-methylpyrazine and 2,3,5-trimethylpyrazine, suggesting that the liver might utilize a variety of understudied sensory receptors to maintain homeostatic functions.^53^ Understanding the molecular recognition of alkylpyrazines to OR5K1 may lay the basis for ligand design campaigns and contribute to characterizing the role of this receptor. Here we report two additional agonists relevant to determining the structure-activity relationship profile of OR5K1 ligands and we investigated the interaction of the set of identified agonists within the binding site of OR5K1. To rationalize the effect of ligand substituents in the receptor binding site context, we determined functional data for the most potent compound on two mutants, L104^3.32^ and L255^6.51^. Both ligand information and mutagenesis data guided the model refinement process.

## RESULTS AND DISCUSSION

### OR5K1 agonists

Pyrazines are known for contributing greatly to the aroma of roasted foods,^54–56^ but are also renowned as semiochemicals,^57–61^ compounds that transfer chemical cues between individuals of the same and/or different species, most often eliciting a standardized behavior.^62^ Recently, OR5K1 was characterized as a specialized odorant receptor for the detection of pyrazine-based key food odorants and semiochemicals.^52^ The most potent compound against OR5K1 is compound **1** (2,3-diethyl-5-methylpyrazine, EC_50_ = 10.29 μM). Compounds tested against OR5K1 include molecules with shorter or missing aliphatic chains to the pyrazine moiety (compounds **4**, **6**, **7**, **12**). We also know that the pyrazine itself does not activate this receptor.^52^ Therefore, the activity of OR5K1 molecules is supposed to rely on the presence and position of the aliphatic chains (Table 1). Interestingly, in the screening of pyrazines, the mixture of isomers 2-ethyl-3,5(6)-dimethylpyrazine was found to activate OR5K1 with an EC_50_ of 21.18 μM.^52^ In this work, we isolated the mixture and tested the individual isomers against OR5K1. We found that 2-ethyl-3,6-dimethylpyrazine (compound **2**) has an EC_50_ of 14.85 μM, while 2-ethyl-3,5-dimethylpyrazine (compound **13**) could not be measured to saturation with the concentration range available. This provides precise information on the contribution of the ethyl groups attached to the pyrazine ring.

**Table 1.**
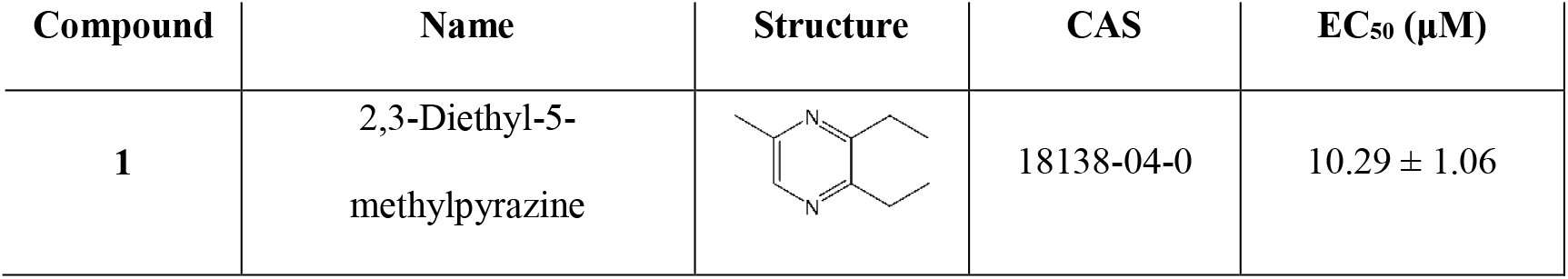
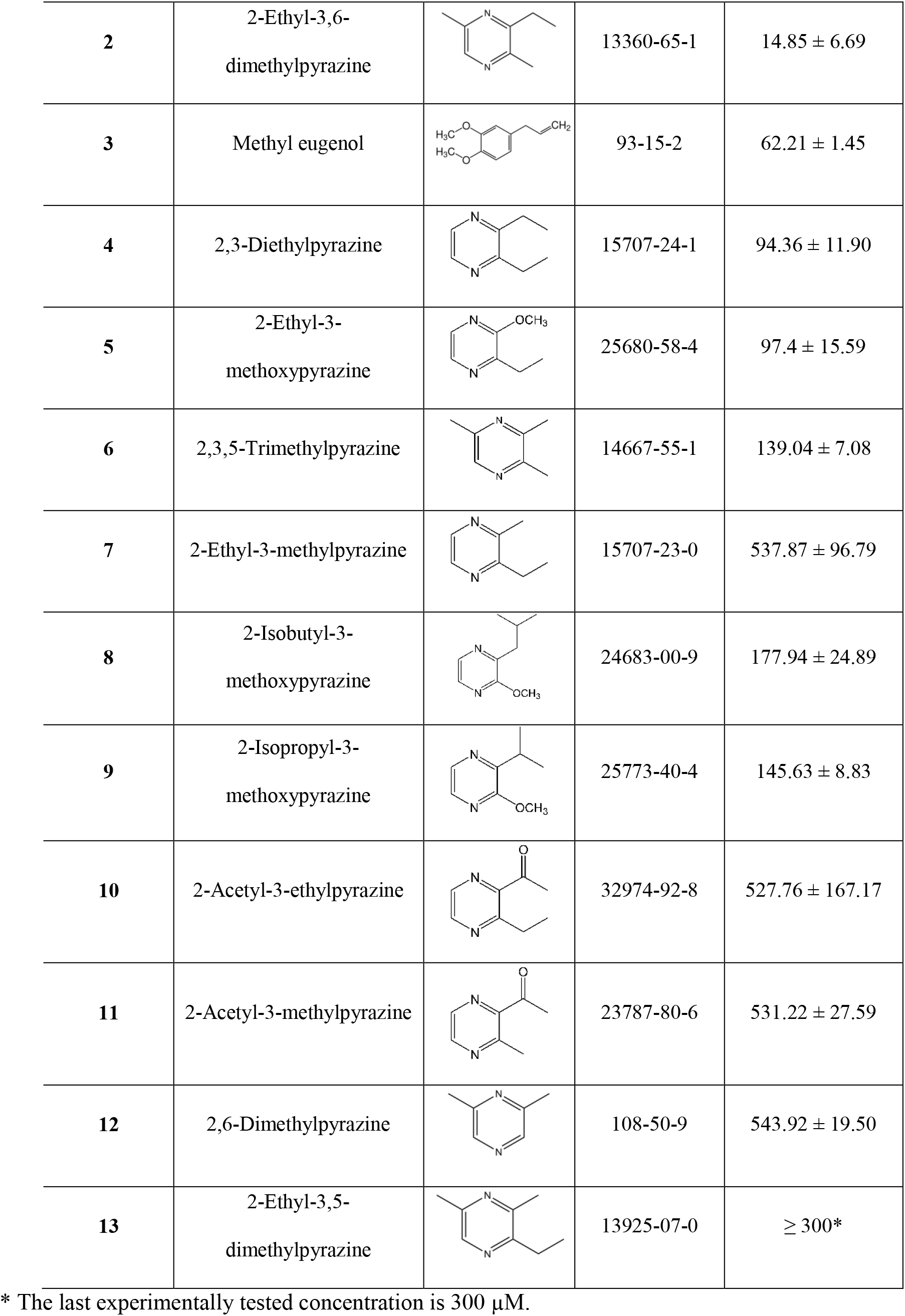
OR5K1 agonists and EC_50_ values. Data for compounds **1**, **3-12** are retrieved from literature,^52^ while data for compounds **2** and **13** were tested in this work (concentration-response curves are reported in Figure S1).

| Compound | Name | Structure | CAS | $EC_{50}$ ($\mu M$) |
| --- | --- | --- | --- | --- |
| **1** | 2,3-Diethyl-5-methylpyrazine | | 18138-04-0 | $10.29 \pm 1.06$ |
| 2 | 2-Ethyl-3,6-dimethylpyrazine | | 13360-65-1 | $14.85 \pm 6.69$ |
| 3 | Methyl eugenol | | 93-15-2 | $62.21 \pm 1.45$ |
| 4 | 2,3-Diethylpyrazine | | 15707-24-1 | $94.36 \pm 11.90$ |
| 5 | 2-Ethyl-3-methoxypyrazine | | 25680-58-4 | $97.4 \pm 15.59$ |
| 6 | 2,3,5-Trimethylpyrazine | | 14667-55-1 | $139.04 \pm 7.08$ |
| 7 | 2-Ethyl-3-methylpyrazine | | 15707-23-0 | $537.87 \pm 96.79$ |
| 8 | 2-Isobutyl-3-methoxypyrazine | | 24683-00-9 | $177.94 \pm 24.89$ |
| 9 | 2-Isopropyl-3-methoxypyrazine | | 25773-40-4 | $145.63 \pm 8.83$ |
| 10 | 2-Acetyl-3-ethylpyrazine | | 32974-92-8 | $527.76 \pm 167.17$ |
| 11 | 2-Acetyl-3-methylpyrazine | | 23787-80-6 | $531.22 \pm 27.59$ |
| 12 | 2,6-Dimethylpyrazine | | 108-50-9 | $543.92 \pm 19.50$ |
| 13 | 2-Ethyl-3,5-dimethylpyrazine | | 13925-07-0 | $\geq 300^*$ |

\* The last experimentally tested concentration is 300 μM.

### OR5K1 structure prediction

ORs and chemosensory GPCRs share low sequence similarity (below 20%) with experimentally solved GPCRs.^20, 63^ The accuracy of 3D structures obtained by homology modeling is highly dependent on the templates. Good models of membrane proteins can be obtained for template sequence identities higher than 30%.^64^ A multi-template homology modeling approach has been used for successfully modeling different ORs, including OR51E1 and OR7D4.^23, 65^ In this approach, conserved motifs were used to guide the sequence alignment of odorant receptors. To obtain a model that could be compared to OR models previously described in literature,^23, 65^ bovine Rhodopsin (bRho), human β2-adrenergic (hβ2AR), human Adenosine-2A (hA2A), and human Chemokine-4 (hCXCR4) receptors were used as templates.^21^ OR5K1 shares 15-19% sequence identity with these templates (Figure S2). Considering that we aimed to use the model to investigate the binding modes of agonists, we built the 3D structure of OR5K1 using bRho, hβ2AR, and hA2A in their active state, while hCXCR4 is only available in its inactive state (the built model is shown in orange in Figure 1).^39^

**Figure 1:**
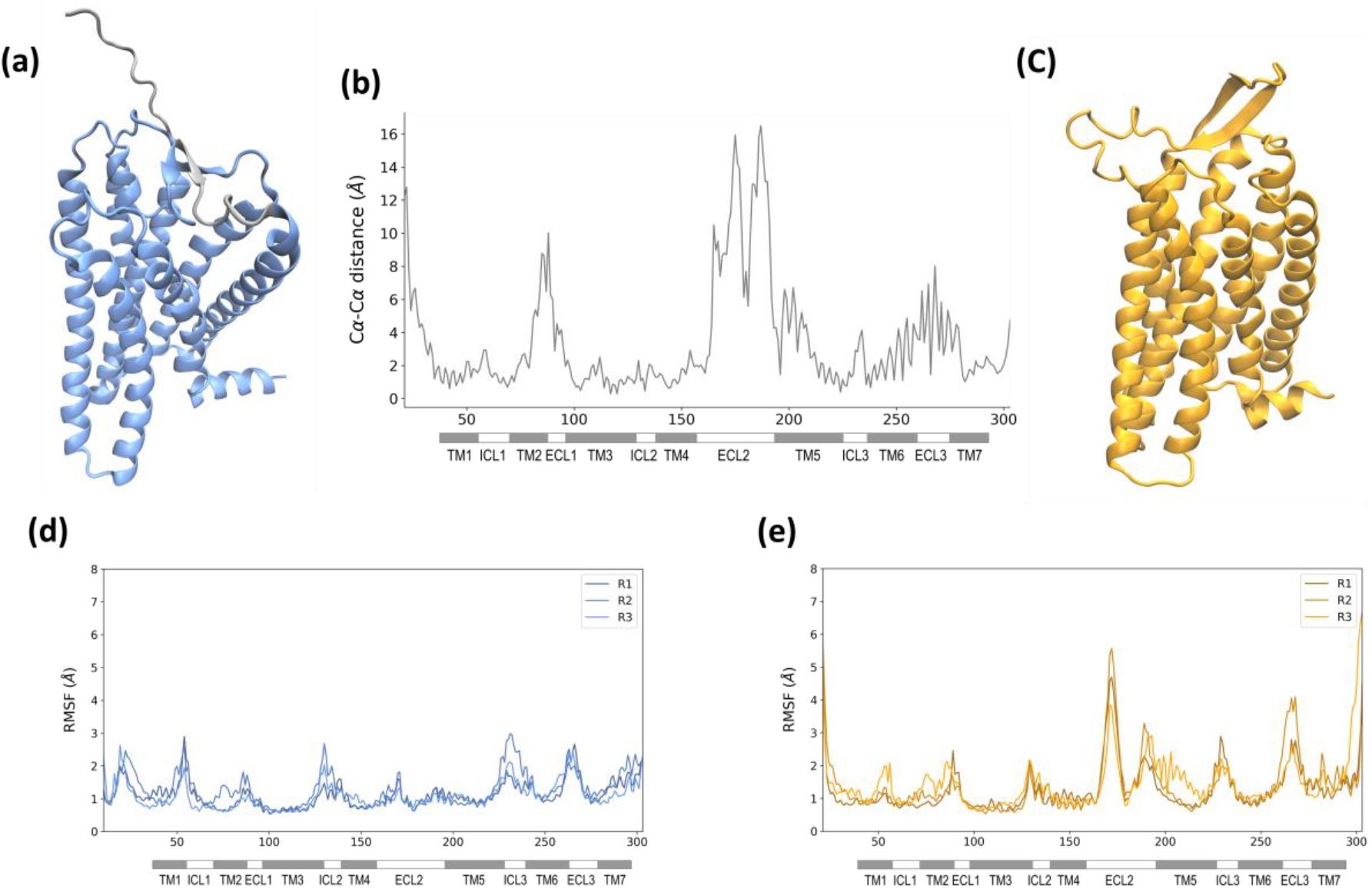
3D representations of AF2 model (a) and HM (c). The N-terminus of the AF2 model is shown in grey. Cα-Cα distances per residue between the two models (b). RMSF plots of Cα atoms through MD replicas (R1, R2, R3) for AF2 model (d) and HM (e).

We then downloaded the Alphafold 2 (AF2) structure of OR5K1 (https://alphafold.ebi.ac.uk/entry/Q8NHB7, the model colored in blue in Figure 1) to compare with the homology model (HM). Except for the N-terminus and the ECL3, the per-residue confidence score (average predicted local distance difference test, pLDDT) of all regions of the model is >90 (very high) or between 70 and 90 (confident) (Figure S3). The OR5K1 AF2 model is also among the high-confidence AF2 GPCR models, as assessed by the per-model pLDDT_80_ score, which was suggested as a potential criterion to assess the quality of AF2 models for structure-based virtual screening.^66^

We calculated the GPCR activation index of the AF2 and HM models using the A100 tool,^67^ confirming that the HM is in its active state with an activation index of 68.46, but AF2 is an inactive model with an activation index of −21.30. In the AF2 database, the activation state is not specifically taken into consideration, and 63% of class A GPCRs are modeled in the inactive state.^68–69^ The different conformational states affect the differences in the 3D structural architecture and the binding site conformations.

AF2 and HM models have a Root Mean Square Deviation (RMSD) of Carbons alpha (Cα) of 4.76 Å. To get a measure of the differences between the two models in the GPCR domains, we calculated and plotted the distances between Cα of the two models for all residues (Figure 1b). The ECL1 and ECL2 are the most different regions in the two models. Also, the two models present an average Cα-Cα distance higher than 4 Å for TM5 residues and in residues 240-270, including the end of TM6, ECL3, and the beginning of TM7 (Figure 1b). TM5 is closer to the orthosteric binding site in the HM than in the AF2 model, and this is also due to the different folding of the ECL2. The secondary structure of the terminal region of TM6 is not well defined in the AF2, this portion is classified with local prediction confidence pLDDT between 70 and 90 for the helix part and lower than 70 for the ECL3 part (Figure S3).

Differences in some regions of the models are also consequent of the different sequence alignments that led to the two models, e.g. we observed a shift of one position in TM7.

We further explored structural differences between the two models with short runs (100ns x 3 replicas) of Molecular Dynamics (MD) simulations. As shown in the Root Mean Square Fluctuation (RMSF) plots (Figure 1d), the AF2 model is rather stable, while we can observe higher fluctuations in the HM, especially in the region of the ECL2.

### The ECL2 of OR5K1

As mentioned above, the ECL2 folding is the most evident difference between the two models. The ECL2 is the largest and most structurally diverse extracellular loop of GPCRs,^70^ and those of ORs are among the longest ECL2 in class A GPCRs.^71^ Loop modeling is highly challenging when sequence length reaches the size of the OR ECL2.^72–74^ A template selection based on sequence identity is rather difficult considering the high sequence and shape variability. In Figure S4a, we report the length of ECL2 segments for OR5K1 and experimental class A GPCRs. The templates chosen for OR5K1 modeling have an ECL2 that is much shorter than the ECL2 of OR5K1. We remodeled this region using templates with higher similarities in terms of length and sequence composition (Figures S2-S4). Specifically, the ECL2 of NPY2 and CCK1 receptors were the templates for the segment before the conserved Cys^45.50^ (S156^4.57^ - C180^45.50^) and the Apelin receptor for the segment after the Cys^45.50^ (C180^45.50^ - L197^5.37^). Therefore, the HM model has an ECL2 with an anti-parallel β-sheet. Differently, AF2 carries out a β-strand forming a β-sheet with the N-terminus and ends with a small α-helix inside the orthosteric binding site. We have previously analyzed the ECL2 experimental and MD structures of class A GPCRs and identified seven different shapes for this loop, represented by a t-distributed stochastic neighbor embedding (t-SNE) analysis (clusters A-F in Figure 2).^70^ Here, we include in this analysis also HM and AF2 structures. Considering the high fluctuation of the ECL2 of the HM model, we added MD frames only from the AF2 MD simulations. The ECL2 in HM was modeled using templates with cluster B folding, and in the ECL2 space, it falls in this region. Instead, AF2 differs from GPCR ECL2 folds, and groups in a separate region of the ECL2 space (Figure 2, black dots).

**Figure 2:**
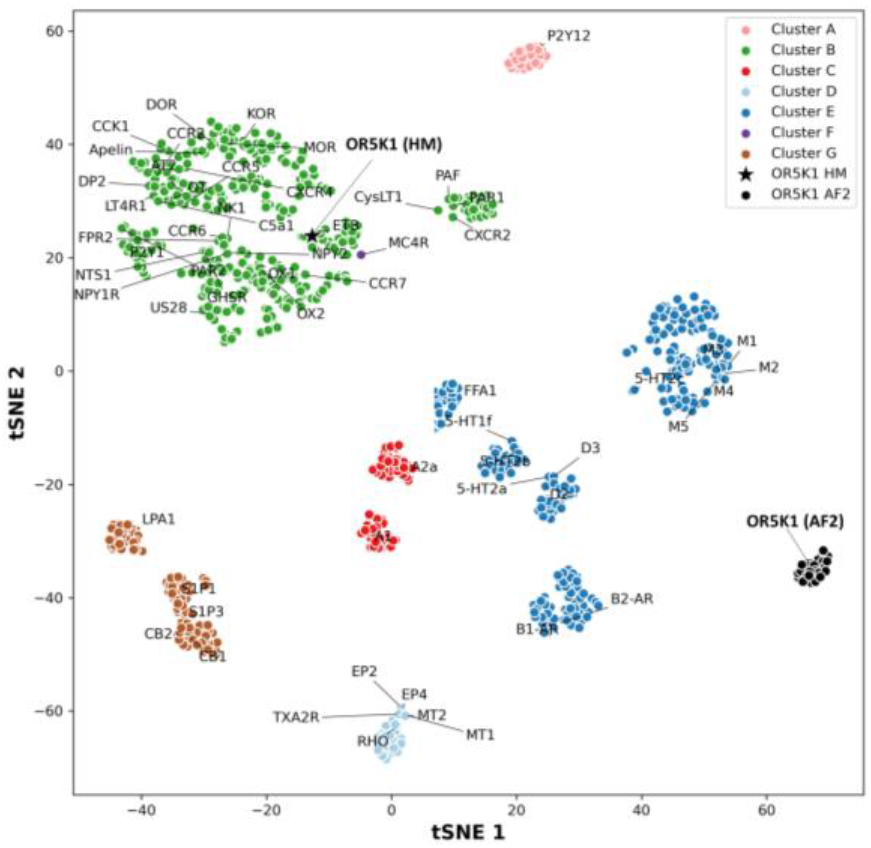
GPCR ECL2 space. In the t-SNE plot, the ECL2 of OR5K1 models are shown in black (HM as a star, MD frames from AF2 model as dots), and experimentally solved GPCRs are colored as pink, dark green, red, light blue, dark blue, violet, and brown, for clusters A, B, C, D, E, F, G, respectively. Data of ECL2 GPCR clusters from Nicoli et al. 2022.^70^

### Binding site sampling

To assess the predictive ability of the HM and AF2 models, we performed molecular docking calculations of known ligands as actives (13 compounds, Table 1) and of all the compounds that did not elicit receptor response with a defined chirality (131 compounds, the complete list with SMILES is available at https://github.com/dipizio/OR5K1_binding_site) as inactives, and we then evaluated the performance of each model through Receiver Operating Characteristic (ROC) analysis.^75–76^ The Area Under the Curve (AUC) values are similar for HM (0.67) and AF2 (0.68), and the enrichment factor in the top 15% of the sorted screened molecules (EF_15%_) is very low in both cases, 0.11 and 0.24 for HM and AF2, respectively (EF_15% max_ = 1.63) (Figure S5). The AF2 model is not able to dock the most potent agonists in our set. The only highly ranked agonist in both HM and AF2 models is compound **9** (EC_50_ = 527.76 μM), with docking scores of −5.68 and −4.91 kcal/mol, respectively. As expected, HM and AF2 models have different residue arrangements in the orthosteric binding site. And in this particular case, the orthosteric binding site of AF2 is not accessible, and the extracellular ligand pocket is located between TM5 and TM6 and extends toward the membrane bilayer (Figure S5). AF2 models are indeed built as *apo* structures and the modeling of binding pocket conformations is not guided by explicit ligand information. Therefore, although evidence of the excellent performance of AF2, especially when no good templates are available, AF2 models might not be *ready-to-use* for structure-based studies.^69, 77–81^

To optimize the binding site, we need to sample the conformational space allowing for residue flexibility. For this purpose, we used induced-fit docking (IFD), an approach that was already applied to GPCR models, including ORs.^82–85^ Using this technique, we can select specific residues to be sampled excluding regions of uncertain modeling. On the contrary, MD simulations can optimize the binding site while taking into account the entire structure flexibility, and this is highly affected by the quality of the model.^86–87^ We performed IFD simulations with the most active compounds (compound **1**) for both AF2 and HM, allowing the binding site side chains to be flexible. 44 models were generated starting from the AF2 model and 57 from HM. The ROC curves of these models show an improvement in their performance, the best models have AUC values of 0.81 and 0.85, and EF_15%_ of 0.24 and 0.50 for AF2 and HM, respectively (Figure S6). The binding modes of compound **1** in the best models of AF2 and HM are different but the ligand is now located in the core of the orthosteric binding site in both models (Figure S6). Interestingly, we noticed that two leucine residues, L104^3.32^ and L255^6.51^, are predicted to be in the binding pocket by both models (Figure S6).

### Key residues for OR5K1 activity

L104^3.32^ is conserved in 10.6% of human ORs, while L255^6.51^ in 15.5% of ORs (Figure S8), but both are strongly conserved in OR5K1 orthologs across species (Figure S9). L104^3.32^ is conserved in 98% of OR5K1 orthologs investigated across 51 species, except for the receptor of the new world monkey *Aotus nancymaae* (XP_012332612.1), where a rather conservative amino acid exchange replaced the leucine at position 104 by isoleucine (Figure S9, Table S5). Similarly, L255^6.51^ of OR5K1 is conserved in 96% of all orthologs investigated, except for the receptors of *Aotus nancymaae, Loxodonta africana* (African elephant, XP_003418985.1), and *Urocitellus parryii* (Arctic ground squirrel, XP_026258216.1). In all three orthologs and in the human paralog OR5K2, again, a rather conservative amino acid exchange replaced the leucine at position 255 with isoleucine (Figure S7, Table S5). Single nucleotide missense variations have been reported for both amino acid positions, L104^3.32^I (rs777947557) and L255^6.51^F (rs1032366530) in human OR5K1, albeit with frequencies way below 0.01. Moreover, both positions L104^3.32^ and L255^6.51^ are part of a set of 22 amino acids that have been suggested previously to constitute a generalized odorant binding pocket in ORs.^88^ Both amino acid positions have been identified also experimentally as odorant interaction partners in different receptors by several independent studies.^24, 36, 65, 89–94^ Therefore, these leucine residues are likely to play a relevant role in the ligand recognition of OR5K1 agonists. We mutated these residues to alanine (L104^3.32^A, L255^6.51^A) and found that there is a shift in EC_50_ values for both mutants when stimulated with compound **1**: EC_50_ of 525.28 ± 92.28 μM for L104^3.32^A and EC_50_ of 478.36 ± 185.10 μM for OR5K1 L255^6.51^A (Figure 3a). The effect of these two leucine residues on OR5K1 activation has been confirmed also for the 2-ethyl-3,5(6)-dimethylpyrazine (Figure S11a).

**Figure 3:**
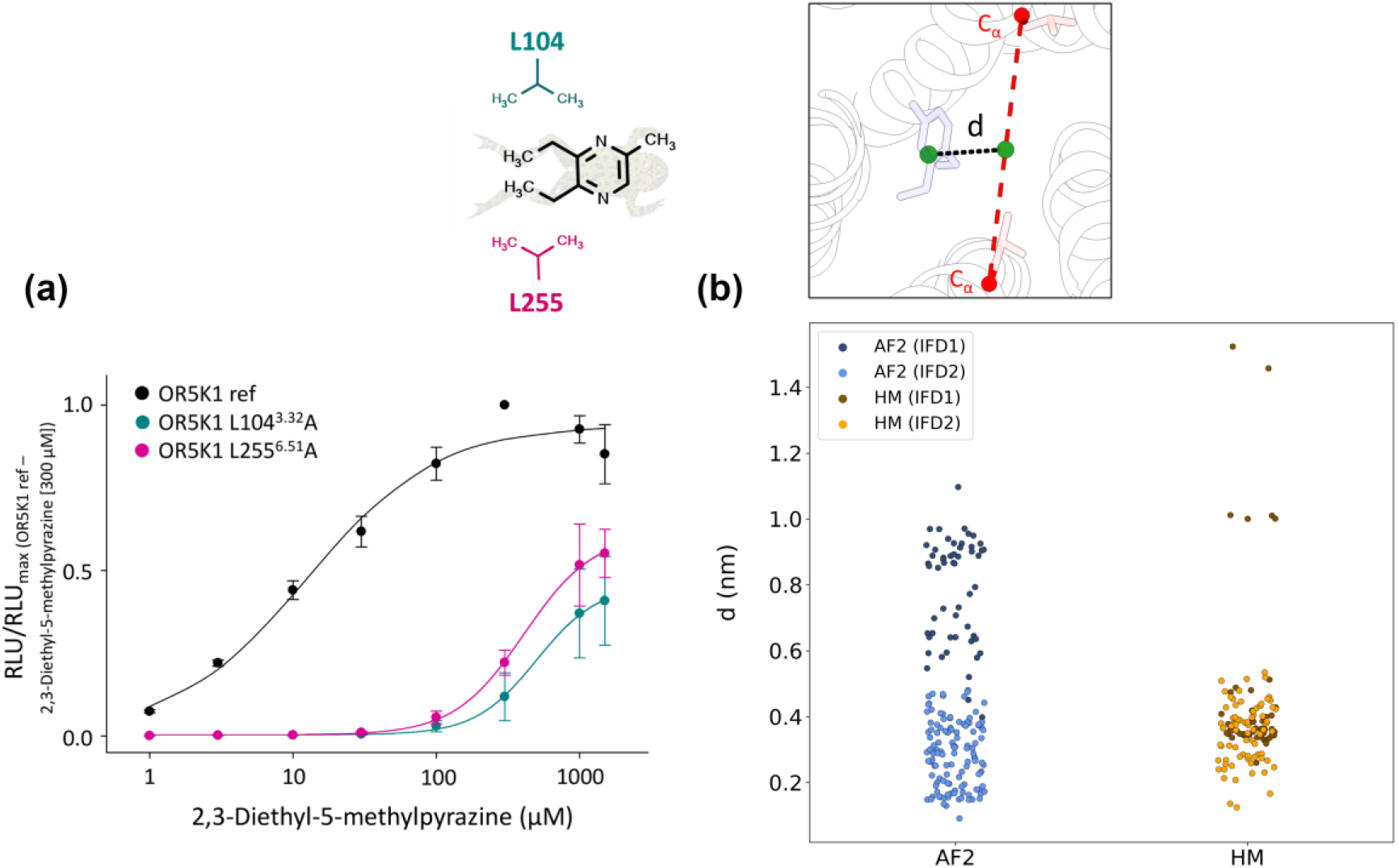
**(a)** Concentration-response relations of compound **1** (2,3-diethyl-5-methylpyrazine) on OR5K1 (black), OR5K1 L104^3.32^A (turquoise), and OR5K1 L255^6.51^A (pink). Data were mock control-subtracted, normalized to the response of OR5K1 ref to 2,3-diethyl-5-methylpyrazine (300 μM) and displayed as mean ± SD (n = 4). RLU = relative luminescence units. **(b)** Distance between the ligand centroid and the center between L104^3.32^ and L255^6.51^ alpha carbons in the first and second IFD simulation rounds (IFD1 and IFD2).

### OR5K1 model refinement

Monitoring the distance between the centroid of the ligand and the center between the Cα atoms of the two leucine residues on the poses obtained with the first round of IFD simulations, we observed that, while for the HM, this distance reaches 0.2 nm, for the AF2 model it is above 0.4 nm (Figure 3b). To improve the conformational rearrangement around the ligand, we performed a second round of IFD simulations, allowing the flexibility of the binding site side chains around compound **1**. With the second round of simulations, there is a better sampling for HM conformations and an enrichment of poses in close contact with L104^3.32^ and L255^6.51^ for the AF2 model (Figure 3b).

Then we analyzed all the poses where the ligand is close to L104^3.32^ and L255^6.51^ (with a distance below 0.4 nm): 106 structures for AF2 (1 from the first round of IFD and 105 from the second round) and 110 for HM (39 from the first round of IFD and 71 from the second round). We clustered the complexes into 31 and 34 possible binding poses for AF2 and HM, respectively. The distribution of the clusters is reported in Figure S7. Among all the potential binding modes, 6 models from the refinement of AF2 model and 12 structures from the refinement of HM have an AUC higher than 0.8 (Table S1). These may be considered the most predictive binding site conformations and were submitted to a third round of IFD simulations for the extensive sampling of the conformational space of L104^3.32^ and L255^6.51^. This generates 555 structures from the model refined from HM and 431 structures from the model refined from AF2 with AUC greater than 0.8 and a distance between the ligand centroid and the center between L104^3.32^ and L255^6.51^ alpha carbons lower than 0.4 nm. The three different rounds of IFD simulations aim to progressively decrease the number of flexible residues (Figure 4a), that is extracellular domain residues (see Methods for the list) in IFD1, residues close to the ligand in IFD2, and L104^3.32^ and L255^6.51^ in IFD3. In Figure 4b, we plot the RMSD values of the binding site with respect to the starting models vs. the AUC values to give an idea of how the structures changed with IFD simulations. The RMSD ranges are defined by the binding site rearrangement sampled with the first round of simulations, but by decreasing the flexible residues in the selection, the conformational space could be more accurately sampled, allowing us to improve the performance (Figure S12). The distribution of AUC and EF values in the three rounds of simulations is visualized in Figure S12. In Figure 4c, we report the poses with the highest AUC values after each IDF round, to show how the binding site is rearranged. We noticed that F202^5.42^ and F256^6.52^ point to the binding site in the starting structure of HM, but not after the IFD optimization, nor for the refined structures of AF2. We could experimentally confirm that these two positions do not affect OR5K1 activation by 2-ethyl-3,5(6)-dimethylpyrazine (Figure S11b).

**Figure 4.**
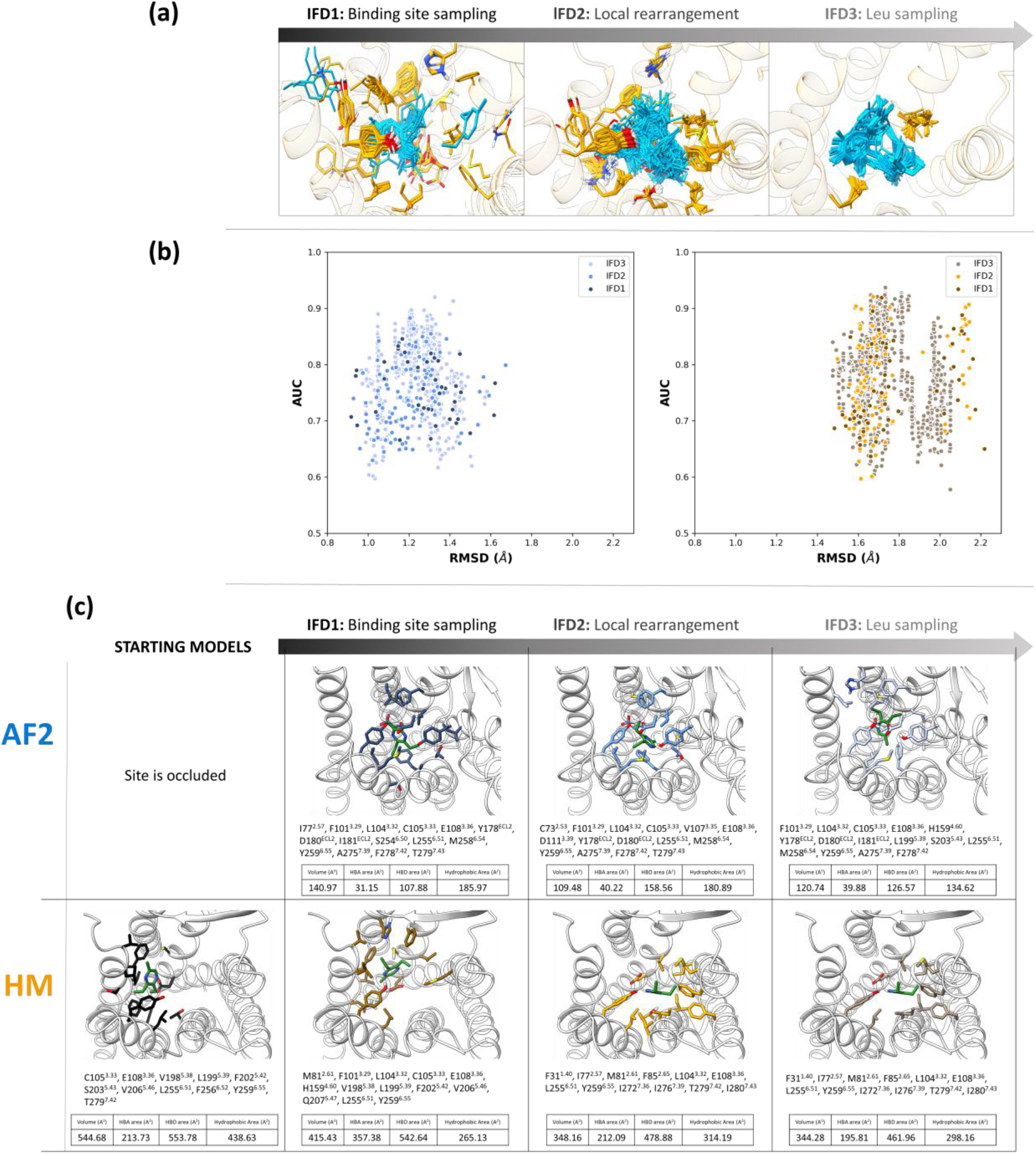
(a) Schematic representation of the three rounds of IFD, showing the decreasing number of selected flexible residues and sampling of ligand binding poses. (b) Plots of RMSD of binding site residues Cα with respect to the starting structures vs. AUC values for all complexes obtained starting from AF2 (blue shades) and HM (orange shades). (c) Binding site residues (at 4 Å from compound **1**) of starting models and best performing models after IFD refinement.

In IFD3, only two residues are sampled. Interestingly, despite the high similarity of structures generated from IFD3 from each system, we could still appreciate different sampled binding modes (37 clusters from HM and 30 clusters from AF2, Figure S13) and performance variability (Figure 4b). The best-performing IFD3 structures for each cluster are available at https://github.com/dipizio/OR5K1 binding site. The binding poses shown in Figure 5 were selected considering the performance, the shape of the ROC curves and the contribution to the binding of L104^3.32^ and L255^6.51^. The ligand in both models is oriented in a similar position and interacts with L104^3.32^ and L255^6.51^. L104^3.32^ and L255^6.51^ interact with the aliphatic chains attached to the pyrazine moiety and might play a relevant role in ligand selectivity. We also performed docking simulations of compound **1** against L104^3.32^A and L255^6.51^A mutant models using the AF2 and HM structures in Figure 5 and we observed in all cases a drop in docking scores (−6.58 and −6.14 kcal/mol for L104^3.32^A and L255^6.51^A mutant AF2 models and −5.58 and −6.02 kcal/mol for L104^3.32^A and L255^6.51^A mutant AF2 models; docking scores obtained with wt models are −7.15 and −6.56 kcal/mol for AF2 and HM, respectively). Therefore, both models seem to be able to capture most differences in activity related to small structural differences either at the ligand or receptor side.

**Figure 5.**
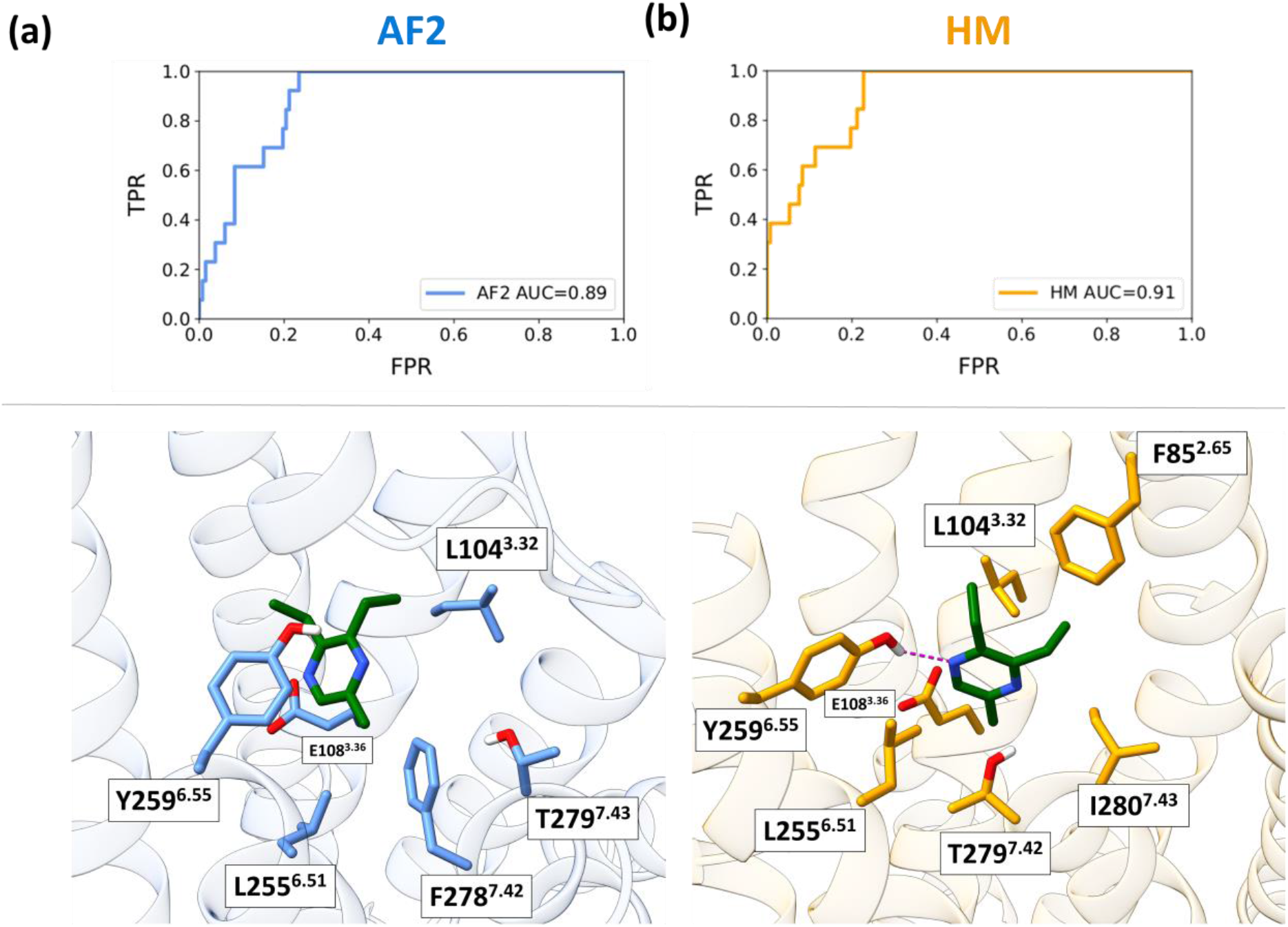
**(a)** ROC curves and **(b)** binding modes of compound **1** into the OR5K1 binding site of the best AF2 and HM models obtained after the extensive sampling of the conformational space of L104^3.32^ and L255^6.51^. We show positions that are in common between the two models as stick residues in the binding site. Residue F85^2.65^ is only reported for the HM model because TM2 in the AF2 model is not pointing to the binding site (the Cα atoms of F85 in the two models are 8.85 Å distant).

## CONCLUSIONS

Odorant molecules are typically small organic compounds of less than 300 Da with high-to-moderate hydrophobicity and their binding to ORs is driven by shape complementarity and mostly hydrophobic interactions.^71, 95^ ORs are class A GPCRs for which we do not have experimental structures and that share very low sequence identity with non-sensory GPCRs. The small size of OR modulators and the low resolution of the structure modeling pose a major challenge to the investigation of the molecular recognition mechanisms of this important class of receptors. Most ORs are still orphans and the receptive range of a few ORs has been characterized until now. In this paper, we used mutagenesis and ligand information to sample and select OR5K1 orthosteric binding site conformations. To enrich the set of agonists with data relevant to determining the structure-activity relationship profile, the mixture of isomers 2-ethyl-3,5(6)-dimethylpyrazine was isolated and the individual isomers were tested against OR5K1. We found that 2-ethyl-3,6-dimethylpyrazine (compound **2**) has an EC_50_ of 14.85 μM, whereas 2-ethyl-3,5-dimethylpyrazine (compound **13**) has an EC_50_ higher than 300 μM.

To generate the starting conformation of OR5K1, we used a multi-template homology modeling approach, as previously suggested to be a successful strategy for OR modeling.^20–21, 23, 65^ Moreover, we further refined the ECL2 loop, which we previously identified to be a necessary procedure for low-resolution GPCR modeling.^75, 87, 96^ We also used the AlphaFold 2 model of OR5K1 for our analyses. A major difference between our HM and AF2 models is in the ECL2 folding. The ECL2 predicted by AF2 seems unique, and was found to be rather stable in MD simulations. Moreover, we found a binding site occlusion and a scale in the TM7 backbone that can compromise the applicability of the AF model for structure-based investigations, as observed also in other studies.^69, 80, 97^

We found that the optimization of the binding site was a necessary step for both HM and AF2 models. The refinement process of AF2 model was needed not only to improve the performance, but also to open the orthosteric binding site and allow the docking of agonists. The location of the orthosteric binging site was driven by the selection of flexible residues in IFD1. The starting models obtained from AF2 and HM have different conformations of the TM helices that prevented reaching convergence when sampling only the side chain conformations. As an example, in Figure 4, it is possible to appreciate the difference in the shift of TM7 residues in the two models: position 7.42 is represented by F278 in the model from AF2, but by T279 in the model from HM. Only when the experimental structure will be solved we can assess which models capture better the structural features of OR5K1. However, our work demonstrates that it is possible to build predictive structural models despite their quality.

Through the modeling, we could identify relevant residues for the activity of OR5K1 agonists, namely, L104^3.32^ and L255^6.51^. Increased EC_50_ values were obtained when compound **1** was tested against OR5K1 mutants L104^3.32^A and L255^6.51^A. Interestingly, 3.32 and 6.51 positions are highly conserved in OR5K1 orthologs across 51 species and have an extremely low frequency of SNP-based missense variations according to the 1000 Genomes Project. The support of mutagenesis experiments furnished precious experimental information for model refinement.

In summary, we propose here an iterative experimental-computational workflow that allowed us to explore the conformational space of the OR5K1 binding site and can be used to model the orthosteric binding site of ORs as well as that of GPCRs with low sequence identity to available templates.

## MATERIALS AND METHODS

### Synthesis of 2-ethyl-3,5(6)-dimethylpyrazine

2-ethyl-3,5(6)-dimethylpyrazines were synthesized according to Czerny et al.^98^ by a Grignard-type reaction. Briefly, a solution of ethylmagnesium bromide in tetrahydrofuran (20 mL; 1.0 M; 20 mmol) was placed in a three-necked flask (100 mL) equipped with a reflux condenser, a dropping funnel and an argon inlet. While stirring at 40 °C a small portion of the respective reactant (2.2 g; 20 mmol) solved in 20 mL THF was added dropwise via the dropping funnel. 2,5-dimethylpyrazine was used for the synthesis of 2-ethyl-3,6-dimethylpyrazine and 2,6-isomere was taken as starting material for 2-ethyl-3,5-dimethylpyrazine. After the mixture was refluxed (73°C) the residual 2,5(6)-dimethylpyrazine solution was added over a period of 30 min. The mixture was stirred under refluxed for 2 h, cooled to room temperature, and water (20 mL) was added dropwise. The emulsion was extracted with diethyl ether (3 ×50 mL) and dried over anhydrous Na2SO4. The compounds were purified by means of flash column chromatography. For this purpose, the concentrated extract (1.0 mL) was placed on the top of a water-cooled glass column (33 × 2.5 cm) filled with a slurry of silica gel 60 (with the addition of 7 % water, 40 – 63 μm, Merck, Darmstadt, Germany, # 1.09385.2500) and n-pentane. The target compounds were eluted with n-pentane/diethyl ether (100 ml, 40:60, v/v). The purity of each target compound was analyzed by gas chromatography-mass spectrometry (GC-MS) and nuclear magnetic resonance (NMR). For determining the concentration of each 2-ethyl-3,5(6)-dimethylpyrazine, quantitative NMR (qNMR) was applied. For the NMR experiments, the solvent was distilled off and the residue was solved in CDCl_3_.

2-ethyl-3,5-dimethylpyrazine: MS (EI): *m/z* (%) 135 (100), 136 (M^+^, 81), 42 (18), 108 (17), 107 (15), 56 (12). ^1^H-NMR (CDCl_3_, 400 MHz, 25 °C) *δ* (ppm) 8.15 (s, 1 H, H-C6), 2,80 (q, *J*=7.6, 2H, H-C7), 2.53 (s, 3 H, H-C9/10, 2.49 (s, 3 H, H-C9/10), 1,27 (t, *J*=7.6, 3H, H-C8).

2-ethyl-3,6-dimethylpyrazine: MS (EI): *m/z* (%) 135 (100), 136 (M^+^, 92), 56 (24), 108 (16), 42 (12), 107 (11). ^1^H-NMR (400 MHz, CDCl_3_) *δ* (ppm) 8.20 (s, 1 H, H-C6), 2.81 (q, *J*=7.5, 2H, H-C7), 2.54 (s, 3 H, H-C9/10, 2.49 (s, 3 H, H-C9/10), 1,28 (t, *J*=7.5, 3H, H-C8).

### Nuclear magnetic resonance (NMR)

NMR experiments were performed using an Avance III 400 MHz spectrometer equipped with a BBI probe (Bruker, Rheinstetten, Germany). Topspin software (version 3.2) was used for data acquisition. For structure elucidation the compounds were solved in chloroform-d (CDCl_3_). Chemical shifts were referenced against solvent signal. Quantitative ^1^H-NMR (qNMR) was done according to Frank et al.^99^ For this, an aliquot (600 μL) of the dissolved solutions was analyzed in NMR tubes (5 × 178 mm, Bruker, Faellanden, Switzerland).

### Gas chromatography – mass spectrometry (GC-MS)

Mass spectra of the synthesized pyrazines in the electron ionization mode were recorded using a GC-MS system consisting of a Trace GC Ultra gas chromatograph coupled to a single quadrupole ISQ mass spectrometer (Thermo Fisher Scientific, Dreieich, Germany) as described in more detailed by Porcelli et al.^100^ A DB-1701 coated fused silica capillary column (30 m × 0.25 mm i.d., 0.25 μm film thickness; Agilent, Waldbronn, Germany) was taken for chromatographic separation using the following temperature program: 40°C held for 2 min, then it was raised at 10 °C/min to 230°C (held for 4 min). Mass spectra were acquired at a scan range of 40–300 m/z at ionization energy of 70 eV. The mass spectra were evaluated using Xcalibur 2.0 software (Thermo Fisher Scientific).

### Molecular cloning of OR5K1

The protein-coding region of human OR5K1 (NM_001004736.3) was derived from our previously published OR library.^101^ Amplification was carried out in a touchdown approach using gene-specific primers (Table S2): an initial denaturation (98 °C, 3 min) and ten cycles consisting of denaturation (98 °C, 30 s), annealing (60 °C, decreasing 1 °C per cycle down to 50 °C, 30 s), and extension (72 °C, 1 min), followed by 25 cycles of denaturation (98 °C, 30 s), annealing (50 °C, 30 s), and extension (72 °C, 1 min), finishing with a final extension step in the end (72 °C, 7 min). Insertion of nucleotides into expression vectors was done with T4-DNA ligase (#M1804, Promega, Madison, USA) via EcoRI/NotI (#R6017/#R6435, Promega, Madison, USA) into the expression plasmid pFN210A,^102^ and verified by Sanger sequencing using internal primers (Table S3) (Eurofins Genomics, Ebersberg, Germany).

### PCR-based site-directed mutagenesis

Mutants L104^3.32^ and L255^6.51^ were generated by PCR-based site-directed mutagenesis in two steps. Utilized mutation primers were designed overlapping and are listed in Table S4. Step one PCR was performed in two amplifications, one with the forward vector-internal primer and the reverse mutation-primer, the other with the forward mutation-primer and the reverse vector-internal primer. Amplification was performed with the touchdown approach described above. Both PCR amplicons were then purified and used as template for step two. The two overlapping amplicons were annealed using the following touchdown program: denaturation (98 °C, 3 min), ten cycles containing denaturation (98 °C, 30 s), annealing (start 60 °C, 30 s), and extension (72 °C, 2 min). After this, vector-internal forward and reverse primers were added and 25 further cycles of denaturation (98 °C, 30 s), annealing (50 °C, 30 s), and extension (72 °C, 1 min) were carried out, finishing with a final extension step in the end (72 °C, 7 min). The amplicons were then sub-cloned as described above.

### Cell culture and transient DNA transfection

We utilized HEK-293 cells,^103^ a human embryonic kidney cell-line, as a test cell system for the functional expression of ORs.^104^ Cells were cultivated at 37 °C, 5% CO_2_, and 100% humidity in 4.5 g/L D-glucose containing DMEM with 10% fetal bovine serum, 2 mM L-glutamine, 100 U/mL penicillin, and 100 U/mL streptomycin. Cells were cultured in a 96-well format (Nunclon^™^ Delta Surface, #136102; Thermo Fisher Scientific, Schwerte, Germany) at 12,000 cells/well overnight. Then, cells were transfected utilizing 0.75 μL/well ViaFect^™^ (#E4981, Promega, USA) with the following constructs: 100 ng/well of the respective OR construct, 50 ng/well of chaperone RTP1S,^105^ 50 ng/well of the G protein subunit Gαolf,^106–107^ olfactory G protein subunit Gγ13,^108^ and 50 ng/well of pGloSensor^TM^-22F (Promega, Madison, USA).^109^ The utilized pGloSensor^™^-22F is a genetically engineered luciferase with a cAMP-binding pocket, allowing for measurements of a direct cAMP-dependent luminescence signal. All measurements were mock-controlled, i.e. pFN210A without OR was transfected in parallel.

### Luminescence assay

Concentration-response assays were measured 42 hours post-transfection as described previously.^104^ In short, the supernatant was removed and cells were loaded with a physiological salt buffer (pH 7.5) containing 140 mmol/L NaCl, 10 mmol/L HEPES, 5 mmol/L KCl, 1 mmol/L CaCl2, 10 mmol/L glucose, and 2% of beetle luciferin sodium salt (Promega, Madison, USA). For luminescence measurements, the GloMax^®^ Discover microplate reader (Promega, Madison, USA) was used. After incubation for 50 minutes in the dark, the basal luminescence signal of each well was recorded thrice. Then the odorant, serially diluted in the physiological salt buffer with added Pluronic PE-10500 (BASF, Ludwigshafen, Germany), was applied to the cells and luminescence was measured thrice after ten minutes of incubation time. The final Pluronic PE-10500 concentration on the cells was 0.05%.

### Data analysis of the cAMP-luminescence measurements

The raw luminescence data obtained from the GloMax^®^ Discover microplate reader detection system were analyzed for concentration/response assays by averaging both data points of basal levels and data points after odorant application. For a given luminescence signal, the respective basal level was subtracted and the now corrected data set was normalized to the maximum amplitude of the reference. The data set for the mock control was subtracted and EC_50_ values and curves were derived from fitting the function:

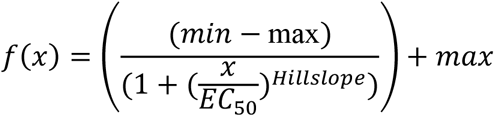

to the data by nonlinear regression (SigmaPlot 14.0, Systat Software).^110^ Data are presented as mean ± SD.

### Flow cytometry

HEK-293 cells were cultivated in 12-well plates with a density of 96,000 cells per well. On the next day, the transfection was performed as described earlier.^111^ For analysis, cells were harvested 42 h post-transfection and stained with the cell-impermeant Halo Tag^®^ Alexa Fluor^®^ 488 Ligand (ex/em= 499/518 nm). Cells were incubated for 30 min at 37°C and 5% CO_2_ in the cell culture incubator. Cells were washed twice with serum free medium prior to FACS analyses (MACSQuant Analyzer, Miltenyi Biotec, Bergisch Gladbach, Germany). A forward- and side-scatter gate was set to exclude dead cells with forward-scatter (FSC: 240V) and side-scatter (SSC: 395V). The FITC signal (B1-channel; HaloTag^®^ Alexa Fluor^®^ 488 Ligand) was detected with 195V. In each case, 10,000 cells were measured. The analysis was performed with the Flowlogic^™^ flow cytometry analysis software (Inivai^™^, Mentone Victoria, Australia). All receptors were measured three times.

### Phylogenetic analysis

NCBI^112^ was used as database for the retrieval of genetic information on *Homo sapiens* (human) odorant receptor genes as well as orthologous receptor genes of OR5K1 (for accession numbers see Table S5). The phylogenetic reconstruction of ORs was performed with QIAGEN CLC Genomics Workbench 21.0 (https://digitalinsights.qiagen.com/) and MEGA X software.^113^ Therefore, in a first step, all sequences were aligned using ClustalW algorithm.^114^ The evolutionary history was inferred using the Neighbor-Joining method ^115^ followed by 500 bootstrap replications.^116^ Scale bar refers to the evolutionary distances, computed using the Poisson correction method.^117^ Evolutionary analyses were conducted in MEGA X.^113^ For rooting the constructed tree, human rhodopsin (NCBI entry: NP_000530.1) was used as an out-group.

### Homology Modeling

Rhodopsin receptor (PDB ID: 4X1H), β2-adrenergic receptor (PDB ID: 6MXT), CXCR4 receptor (PDB ID: 3ODU), and A2A receptor (PDB ID: 2YDV) were used as templates for modeling the 3D structure of OR5K1, following the template selection from de March et al. 2015.^20^ The structures were downloaded from GPCRdb,^118^ and their sequences were aligned to the OR5K1 sequence (residues 20-292) with the Protein Structure Alignment module available in Maestro (Schrödinger Release 2021-3, Maestro, Schrödinger, LLC, New York, NY, 2021). The sequence alignment was then manually adjusted, ensuring that conserved GPCR residues were correctly aligned (Figure S1). OR5K1 shares a sequence identity of 19% with 6MXT.pdb, of 15% with 4X1H.pdb, of 15% with 3ODU.pdb and of 16% with 2YDV.pdb. We modeled the ECL2 region (S157^4.57^- L197^5.37^) using as templates NPY2 (PDB ID: 7DDZ) and CCK1 (PDB ID: 7MBY) for the before-Cys^45.50^ segment, and apelin (PDB ID: 6KNM) for the after-Cys^45.50^ segment (Figures S2 and S3). We also remodeled the region between P81^2.58^ and L104^3.32^ with the NPY2 to ensure the correct orientation of the ECL2 towards TM3 and ECL1, and the formation of the conserved disulfide bridge between C^3.25^ and C^45.50^. 100 homology models were generated using MODELLER v9.23.^119^ Four models were selected based on the DOPE score and visual inspection of the ECL2 and the most predictive model, based on ROC AUC (see the paragraph Molecular Docking) was chosen for the following analysis.

### Protein preparation and binding site analysis

OR5K1 AF2 model was downloaded from the AlphaFold 2 database (https://alphafold.ebi.ac.uk/entry/Q8NHB7). OR5K1 AF2 and HM were superimposed through the Protein Structure Alignment module available in Maestro (Schrödinger Release 2021-3, Maestro, Schrödinger, LLC, New York, NY, 2021). RMSD values were calculated with visual molecular dynamics (VMD).^120^ Hydrogen atoms and side chains of both models were optimized with the Protein Preparation Wizard tool at physiological pH (Schrödinger Release 2021-3, Maestro, Schrödinger, LLC, New York, NY, 2021). Histidine residues 56, 159 and 176 were protonated on the epsilon nitrogen, while all others were protonated on the delta nitrogen. Ramachandran plots were generated to verify the reliability of the backbone dihedral angles of amino acid residues in the models. The A100 tool was used to investigate the activation state of the models.^67^

### Molecular Dynamics simulations

Homolwat webserver (https://alf06.uab.es/homolwat/)^121^ was used to add water molecules within the receptor structures, applying settings described in the GPCRmd protocol.^122^ The prepared structures were then embedded into a 1-palmitoyl-2oleyl-sn-glycerol-3-phospho-choline (POPC) square bilayer of 85Å x 85Å through an insertion method by using HTMD (Accelera, version 2.0.8).^123–124^ The membrane bilayer were previously prepared with VMD Membrane Builder plugin 1.1.

The orientation of the prepared structures within the membrane bilayer were obtained from the coordinates of the β2 adrenergic receptor (PDB ID: 6MXT), as deposited in the Orientations of Proteins in Membranes (OPM) database.^125^ Overlapping lipids were removed upon protein insertion and TIP3P water molecules were added at 15Å from protein atoms by using VMD Solvate plugin 1.5. Finally, the systems were neutralized by Na+/Cl-to reach a final physiological concentration of 0.154 M by using VMD Autonize plugin 1.3.

MD simulations with periodic boundary conditions were carried out with ACEMD^126^ (Acellera, version 3.5.1) using the CHARMM36 force field^127^. The systems were equilibrated through a 3500 conjugate gradient step minimization to reduced clashed between protein and lipid/water atoms, followed by 25 ns of MD simulation in the isothermal–isobaric conditions (NPT ensemble), employing an integration step of 2 fs. Initial constrains were gradually reduced in a three step procedure: positional constrains of 5 kcal mol^−1^ Å^−2^ on lipid phosphorous atoms in the first 5 ns, and positional constrains of 5 kcal mol^−1^ Å^−2^ on protein atoms for the first 15ns; then in the second stage positional constraint were applied only to the protein Cα atoms for other additional 5ns. In the last equilibration stage of 5ns, no restraints were applied. During the equilibration, the temperature was maintained at 310 K using a Langevin thermostat with low damping constant of 1 ps-1, and the pressure was maintained at 1.01325 atm using a Montecarlo barostat. The M-SHAKE algorithm^128^ was used to constrain the bond lengths involving hydrogen atoms. The cutoff distance of 9.0 Å was set for long-term interactions, and 7.5 Å for the switching function. Long-range Columbic interactions were handled using the particle mesh Ewald summation method^129^ (PME) with grid size rounded to the approximate integer value of cell wall dimensions. A non-bonded cutoff distance of 9 Å with a switching distance of 7.5 Å was used.

Equilibrated systems were then subjected to three replicas of 100 ns of unrestrained MD simulation run in the canonical ensemble (NVT) with an integration time step of 4 fs. The temperature was set at 310K, by setting the damping constant at 0.1 ps-1. RMSF plots were computed with an in-house python script based on ProDy (v2.2.0).^130^

### Analysis of ECL2 folds

We used the protocol developed in Nicoli et al. 2022.^70^ AF2 and HM models were superimposed to the kappa-type opioid receptor (KOR, PDB ID: 4DJH) with the Protein Structure Alignment module available in Maestro, Schrödinger (Schrödinger Release 2021-3, Maestro, Schrödinger, LLC, New York, NY, 2021); MD trajectories of AF2 OR5K1 were superimposed using VMD.^120^ Ten representative ECL2 structures were extracted from each replica using the average linkage hierarchical clustering based on backbone volume overlaps (Phase_volCalc and Volume_cluster utilities in Schrödinger).

We added the OR5K1 ECL2 structures (HM and AF2 starting models, and AF2 representative MD frames) to our previous dataset consisting of 60 experimental structures, and 840 MD frames.^70^ Volume overlaps of all ECL2 structures (backbone atoms) were calculated using Phase_volCalc utility from Schrödinger (Schrödinger Release 2021-3, Maestro, Schrödinger, LLC, New York, NY, 2021). Then, pairwise volume overlap values were used to generate a dissimilarity matrix (1-n). The matrix was subjected to a dimensional reduction with t-SNE using the Scikit-learn^131^ (v0.24.2) python module, parameters: angle = 0, perplexity = 25, and 1000 maximum iterations. Visualization of the first two t-SNE components was done with Matplotlib Python library.^132^

### Molecular Docking

The compounds used in the screening by Marcinek et al. were used for the model evaluation.^52^ However, we excluded from this set 54 molecules employed as a mixture of isomers. Indeed, the measured activity of the mixture may not correspond to the activity of the individual stereoisomers (e.g., only one stereoisomer is active) and compromise our validation. Among the subset of molecules with defined stereochemistry, we selected 11 agonists with EC_50_ values below 600 μM and compounds characterized in this work were included in the list of active molecules (Table 1). 131 compounds that did not elicit receptor response were used as inactives (the list of compounds is available at https://github.com/dipizio/OR5K1_binding_site).

3D structures of ligands and inactive molecules were retrieved from PubChem through CAS numbers and prepared for docking through the generation of stereoisomers and protonation states at pH 7.2 ± 0.2 with LigPrep, as implemented in the Schrödinger Small-Molecule Drug Discovery Suite 2021 (LigPrep, Schrödinger, LLC, New York, NY, 2021). Glide Standard Precision (Glide, Schrödinger, LLC, New York, NY, 2021)^133–134^ was used for docking all compounds to the OR5K1 models. The grid box was the centroid of SiteMap grid points for HM and AF2 binding pockets combined together for the models obtained after the first round of IFD, and instead was the centroid of the docked 2,3-diethyl-5-methylpyrazine (compound **1**) for the models obtained after the second round of IFD simulations.

An in-house python script based on Scikit-learn (v0.24.2) package was used for the ROC curve analysis,^131^ and the data were plotted with Matplotlib Python library.^132^ AUC and EF_15%_ of the training library were used to evaluate the performance of each model in discriminating between active and inactive compounds.

The ROC curves were obtained plotting False Positive Rate (FPR) vs. True Positive Rate (TPR). TPR and FPR values are calculated by the following equations:

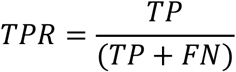

where TP is the number of true positive compounds, and FN is the number of false negative compounds.

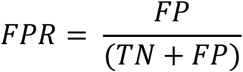

where FP is the number of false positive compounds, and TN is the number of true negative compounds.

EF_15%_ values are calculated by the following equation:

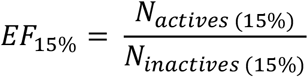

where *N_actives (15%)_* and *N_inactives (15%)_* represent the number of actives and inactives, respectively, in the 15% of ranked screened compounds.

The docking poses of compound **1** within OR5K1 mutants were performed using the *in-place* docking (Glide Standard precision), generating the grid from the centroid of the docked compound. Mutants were generated from the refined models (Figure 5) with the ‘Mutate residue’ tool available in Maestro.

### Induced-fit docking simulations

In the first round of simulations, HM and AF2 starting models were used for IFD simulations using Schrödinger Suite 2021 Induced Fit Docking protocol (Glide, Schrödinger, LLC, New York, NY, 2021; Prime, Schrödinger, LLC, New York, NY, 2021).^135^ 2,3-diethyl-5-methylpyrazine was used as ligand and the flexibility of the side chains at 3 Å from the SiteMap grid points was allowed. The best structures based on AUC values and visual inspection from IFD1 (4 structures after refinement of HM and 7 after refinement of AF2 model) underwent a second round of simulations (IFD2). In the second round of simulations, the residues at 4 Å from the ligand (2,3-diethyl-5-methylpyrazine) were allowed to move. The most predictive structures from IFD2 (Table S1) were submitted to a third round of IFD simulations (IFD 3), in which only the side chains of L104^3.32^ and L255^6.51^ and the ligand were treated as flexible. For an extensive sampling of the leucine residues, we used as the ligand both compounds **1** and **2**.

### Clustering of docking poses

For all poses from IFD1, IFD2, and IDF3 we monitored the distance between the ligand centroid and the center between L104^3.32^ and L255^6.51^ alpha carbons. The centroids and distances were calculated using PLUMED (version 2.7).^136–138^ The docking poses from IDF1 and IDF2 with a distance below 0.4 nm were clustered using the conformer_cluster.py from Schrödinger (https://www.Schrödinger.com/scriptcenter). First, a pair-wise RMSD matrix was calculated for compound **1** and the residues within 7 Å of its centroid (for HM, residues 104, 105, 108, 159, 199, 202, 206, 255, 256, 276, 279, 280; for AF2, residues: 101, 104, 105, 108, 178, 180, 181, 199, 255, 258, 259, 275, 278, 279), and then the complexes were clustered using the hierarchical cluster method (average group linkage). The number of clusters was set to 31 for AF2 and 34 for HM based on the second minimum of the Kelly-Penalty score. Docking poses obtained from IDF3 were filtered by distance (below 0.4 nm), AUC (greater than 0.8) and the conformations of the binding site were clustered using the conformer_cluster.py from Schrödinger. RMSD matrices of best-performing structures from the different clusters were calculated with rmsd.py from Schrödinger (Figure S13).

SiteMap tool (Schrödinger Release 2021-3: SiteMap, Schrödinger, LLC, New York, NY, 2021) was used to characterize the binding cavities of the starting HM and AF2 models, and the best performance models after IFD1, IFD2, and IFD3.

ChimeraX (v1.3) was used to render the protein images.^139^

## DATA AND SOFTWARE AVAILABILITY

The dataset of OR5K1 ligands, starting and refined OR5K1 3D structure models can be downloaded from https://github.com/dipizio/OR5K1_binding_site. MD trajectories are available at https://doi.org/10.5281/zenodo.7182231.

### Supporting Information

Concentration-response relations and activity data of 2-ethyl-3,6-dimethylpyrazine, 2-ethyl-3,5/6-dimethylpyrazine, and 2-ethyl-3,5-dimethylpyrazine on OR5K1 and mutants; sequences of olfactory receptor genes investigated, sequence alignments and phylogenetic analyses; binding site representations, RMSD matrices, ROC analyses, AUC and EF values; clustering results.

## AUTHOR INFORMATION

### Authors

Alessandro Nicoli - Molecular Modeling Group, Leibniz Institute for Food Systems Biology at the Technical University of Munich, 85354 Freising, Germany;

Franziska Haag - Taste and Odor Systems Reception Group, Leibniz Institute for Food Systems Biology at the Technical University of Munich, 85354 Freising, Germany;

Patrick Marcinek - Taste and Odor Systems Reception Group, Leibniz Institute for Food Systems Biology at the Technical University of Munich, 85354 Freising, Germany;

Ruiming He - Molecular Modeling Group, Leibniz Institute for Food Systems Biology at the Technical University of Munich, 85354 Freising, Germany; Department of Chemistry, Technical University of Munich, 85748 Garching

Johanna Kreißl – Analytical Technologies, Leibniz Institute for Food Systems Biology at the Technical University of Munich, 85354 Freising, Germany;

Jörg Stein - Food Metabolome Chemistry Group, Leibniz Institute for Food Systems Biology at the Technical University of Munich, 85354 Freising, Germany

Alessandro Marchetto - Computational Biomedicine group, Institute for Advanced Simulations (IAS)-5/Institute for Neuroscience and Medicine (INM)-9, Forschungszentrum Jülich, 52428 Jülich, Germany; Faculty of Mathematics, Computer Science and Natural Sciences, RWTH Aachen University, 52062 Aachen, Germany;

Andreas Dunkel - Integrative Food Systems Analysis Group, Leibniz Institute for Food Systems Biology at the Technical University of Munich, 85354 Freising, Germany;

Thomas Hofmann - Chair of Food Chemistry and Molecular Sensory Science, Technical University of Munich, 85354 Freising, Germany;

### Author Contributions

The manuscript was written through the contributions of all authors. All authors have approved the final version of the manuscript.

## ACKNOWLEDGMENTS

The authors thank Claire A. de March (Duke University Medical Center) for insightful discussions on OR structures and modeling, and Alexandra Steuer (Leibniz Institute for Food Systems Biology at the Technical University of Munich) and Matteo Pavan (University of Padua) for the critical reading of the manuscript. AN and ADP are members of the COST Actions CA18133, the European Research Network on Signal Transduction (https://ernest-gpcr.eu) and CA18202, the Network for Equilibria and Chemical Thermodynamics Advanced Research (https://www.cost-nectar.eu/). ADP research is supported by the German Research Foundation (PI 1672/3-1).

---

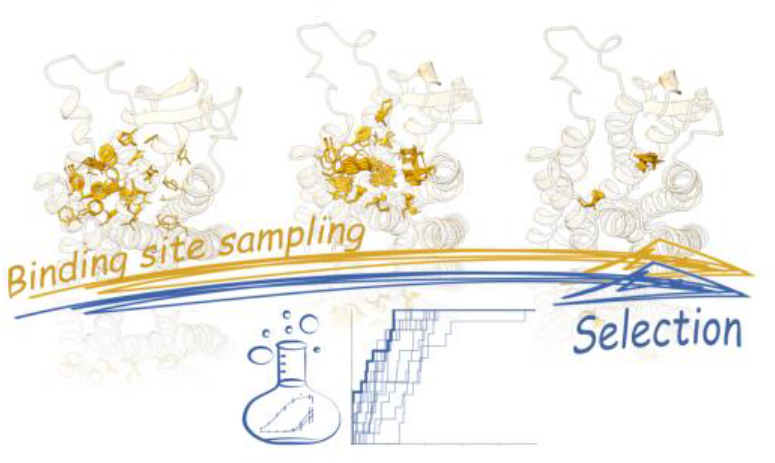

## REFERENCES

(1) Hall, R. A.; Premont, R. T.; Lefkowitz, R. J., Heptahelical receptor signaling: beyond the G protein paradigm. J Cell Biol 1999, 145, 927–32.

(2) Pierce, K. L.; Premont, R. T.; Lefkowitz, R. J., Seven-transmembrane receptors. Nat Rev Mol Cell Biol 2002, 3, 639–50.

(3) Weis, W. I.; Kobilka, B. K., The Molecular Basis of G Protein-Coupled Receptor Activation. Annu Rev Biochem 2018, 87, 897–919.

(4) Fredriksson, R.; Lagerstrom, M. C.; Lundin, L. G.; Schioth, H. B., The G-protein-coupled receptors in the human genome form five main families. Phylogenetic analysis, paralogon groups, and fingerprints. Mol Pharmacol 2003, 63, 1256–72.

(5) Davies, M. N.; Secker, A.; Halling-Brown, M.; Moss, D. S.; Freitas, A. A.; Timmis, J.; Clark, E.; Flower, D. R., GPCRTree: online hierarchical classification of GPCR function. BMC Res Notes 2008, 1, 67.

(6) Nordstrom, K. J.; Sallman Almen, M.; Edstam, M. M.; Fredriksson, R.; Schioth, H. B., Independent HHsearch, Needleman--Wunsch-based, and motif analyses reveal the overall hierarchy for most of the G protein-coupled receptor families. Mol Biol Evol 2011, 28, 2471–80.

(7) Chan, H. C. S.; Li, Y.; Dahoun, T.; Vogel, H.; Yuan, S., New Binding Sites, New Opportunities for GPCR Drug Discovery. Trends Biochem Sci 2019, 44, 312–330.

(8) Congreve, M.; de Graaf, C.; Swain, N. A.; Tate, C. G., Impact of GPCR Structures on Drug Discovery. Cell 2020, 181, 81–91.

(9) Sanchez-Reyes, O. B.; Cooke, A. L. G.; Tranter, D. B.; Rashid, D.; Eilers, M.; Reeves, P. J.; Smith, S. O., G Protein-Coupled Receptors Contain Two Conserved Packing Clusters. Biophys J 2017, 112, 2315–2326.

(10) Hilger, D.; Masureel, M.; Kobilka, B. K., Structure and dynamics of GPCR signaling complexes. Nat Struct Mol Biol 2018, 25, 4–12.

(11) Niimura, Y., Evolutionary dynamics of olfactory receptor genes in chordates: interaction between environments and genomic contents. Hum Genomics 2009, 4, 107–18.

(12) Olender, T.; Waszak, S. M.; Viavant, M.; Khen, M.; Ben-Asher, E.; Reyes, A.; Nativ, N.; Wysocki, C. J.; Ge, D.; Lancet, D., Personal receptor repertoires: olfaction as a model. BMC Genomics 2012, 13, 414.

(13) Buck, L.; Axel, R., A novel multigene family may encode odorant receptors: a molecular basis for odor recognition. Cell 1991, 65, 175–87.

(14) Smith, S. O., Deconstructing the transmembrane core of class A G protein-coupled receptors. Trends Biochem Sci 2021, 46, 1017–1029.

(15) Malnic, B.; Godfrey, P. A.; Buck, L. B., The human olfactory receptor gene family. Proc Natl Acad Sci U S A 2004, 101, 2584–9.

(16) Glusman, G.; Bahar, A.; Sharon, D.; Pilpel, Y.; White, J.; Lancet, D., The olfactory receptor gene superfamily: data mining, classification, and nomenclature. Mamm Genome 2000, 11, 1016–23.

(17) Zhang, X.; Firestein, S., The olfactory receptor gene superfamily of the mouse. Nat Neurosci 2002, 5, 124–33.

(18) Niimura, Y.; Nei, M., Evolutionary dynamics of olfactory receptor genes in fishes and tetrapods. Proc Natl Acad Sci U S A 2005, 102, 6039–44.

(19) Kotthoff, M.; Bauer, J.; Haag, F.; Krautwurst, D., Conserved C-terminal motifs in odorant receptors instruct their cell surface expression and cAMP signaling. FASEB J 2021, 35, e21274.

(20) de March, C. A.; Kim, S. K.; Antonczak, S.; Goddard, W. A., 3rd; Golebiowski, J., G protein-coupled odorant receptors: From sequence to structure. Protein Sci 2015, 24, 1543–8.

(21) de March, C. A.; Yu, Y.; Ni, M. J.; Adipietro, K. A.; Matsunami, H.; Ma, M.; Golebiowski, J., Conserved Residues Control Activation of Mammalian G Protein-Coupled Odorant Receptors. J Am Chem Soc 2015, 137, 8611–8616.

(22) Man, O.; Gilad, Y.; Lancet, D., Prediction of the odorant binding site of olfactory receptor proteins by human-mouse comparisons. Protein Sci 2004, 13, 240–54.

(23) Bushdid, C.; de March, C. A.; Topin, J.; Do, M.; Matsunami, H.; Golebiowski, J., Mammalian class I odorant receptors exhibit a conserved vestibular-binding pocket. Cell Mol Life Sci 2019, 76, 995–1004.

(24) Gelis, L.; Wolf, S.; Hatt, H.; Neuhaus, E. M.; Gerwert, K., Prediction of a ligand-binding niche within a human olfactory receptor by combining site-directed mutagenesis with dynamic homology modeling. Angew Chem Int Ed Engl 2012, 51, 1274–8.

(25) Cong, X.; Ren, W.; Pacalon, J.; Xu, R.; Xu, L.; Li, X.; de March, C. A.; Matsunami, H.; Yu, H.; Yu, Y.; Golebiowski, J., Large-Scale G Protein-Coupled Olfactory Receptor–Ligand Pairing. ACS Central Science 2022, 8, 379–387.

(26) Chen, H.; Dadsetan, S.; Fomina, A. F.; Gong, Q., Expressing exogenous functional odorant receptors in cultured olfactory sensory neurons. Neural development 2008, 3, 22.

(27) Geithe, C.; Noe, F.; Kreissl, J.; Krautwurst, D., The Broadly Tuned Odorant Receptor OR1A1 is Highly Selective for 3-Methyl-2,4-nonanedione, a Key Food Odorant in Aged Wines, Tea, and Other Foods. Chemical senses 2017, 42, 181–193.

(28) Kajiya, K.; Inaki, K.; Tanaka, M.; Haga, T.; Kataoka, H.; Touhara, K., Molecular bases of odor discrimination: Reconstitution of olfactory receptors that recognize overlapping sets of odorants. J Neurosci 2001, 21, 6018–25.

(29) Malnic, B.; Hirono, J.; Sato, T.; Buck, L. B., Combinatorial receptor codes for odors. Cell 1999, 96, 713–23.

(30) Nara, K.; Saraiva, L. R.; Ye, X. L.; Buck, L. B., A Large-Scale Analysis of Odor Coding in the Olfactory Epithelium. Journal of Neuroscience 2011, 31, 9179–9191.

(31) Saito, H.; Chi, Q.; Zhuang, H.; Matsunami, H.; Mainland, J. D., Odor coding by a Mammalian receptor repertoire. Sci Signal 2009, 2, ra9.

(32) Adipietro, K. A.; Mainland, J. D.; Matsunami, H., Functional evolution of mammalian odorant receptors. PLoS Genet 2012, 8, e1002821.

(33) Mainland, J. D.; Li, Y. R.; Zhou, T.; Liu, W. L.; Matsunami, H., Human olfactory receptor responses to odorants. Scientific data 2015, 2, 150002.

(34) Noe, F.; Polster, J.; Geithe, C.; Kotthoff, M.; Schieberle, P.; Krautwurst, D., OR2M3: A Highly Specific and Narrowly Tuned Human Odorant Receptor for the Sensitive Detection of Onion Key Food Odorant 3-Mercapto-2-methylpentan-1-ol. Chemical senses 2017, 42, 195–210.

(35) Haag, F.; Di Pizio, A.; Krautwurst, D., The key food odorant receptive range of broadly tuned receptor OR2W1. Food Chem 2022, 375, 131680.

(36) Charlier, L.; Topin, J.; Ronin, C.; Kim, S. K.; Goddard, W. A., 3rd; Efremov, R.; Golebiowski, J., How broadly tuned olfactory receptors equally recognize their agonists. Human OR1G1 as a test case. Cell Mol Life Sci 2012, 69, 4205–13.

(37) Jabeen, A.; de March, C. A.; Matsunami, H.; Ranganathan, S., Machine Learning Assisted Approach for Finding Novel High Activity Agonists of Human Ectopic Olfactory Receptors. Int J Mol Sci 2021, 22.

(38) Di Pizio, A.; Behr, J.; Krautwurst, D., Toward the Digitalization of Olfaction. In Reference Module in Neuroscience and Biobehavioral Psychology, 2021.

(39) Ballante, F.; Kooistra, A. J.; Kampen, S.; de Graaf, C.; Carlsson, J., Structure-Based Virtual Screening for Ligands of G Protein-Coupled Receptors: What Can Molecular Docking Do for You? Pharmacol Rev 2021, 73, 527–565.

(40) Block, E., Molecular Basis of Mammalian Odor Discrimination: A Status Report. J Agric Food Chem 2018, 66, 13346–13366.

(41) Bushdid, C.; de March, C. A.; Fiorucci, S.; Matsunami, H.; Golebiowski, J., Agonists of G-Protein-Coupled Odorant Receptors Are Predicted from Chemical Features. J Phys Chem Lett 2018, 9, 2235–2240.

(42) Yuan, S.; Dahoun, T.; Brugarolas, M.; Pick, H.; Filipek, S.; Vogel, H., Computational modeling of the olfactory receptor Olfr73 suggests a molecular basis for low potency of olfactory receptor-activating compounds. Commun Biol 2019, 2, 141.

(43) Haag, F.; Ahmed, L.; Reiss, K.; Block, E.; Batista, V. S.; Krautwurst, D., Copper-mediated thiol potentiation and mutagenesis-guided modeling suggest a highly conserved copper-binding motif in human OR2M3. Cell Mol Life Sci 2019.

(44) Cong, X.; Ren, W.; Pacalon, J.; Xu, R.; Xu, L.; Li, X.; de March, C. A.; Matsunami, H.; Yu, H.; Yu, Y.; Golebiowski, J., Large-Scale G Protein-Coupled Olfactory Receptor–Ligand Pairing. ACS Central Science 2022.

(45) Hameduh, T.; Haddad, Y.; Adam, V.; Heger, Z., Homology modeling in the time of collective and artificial intelligence. Comput Struct Biotechnol J 2020, 18, 3494–3506.

(46) Akdel, M.; Pires, D. E. V.; Porta Pardo, E.; Jänes, J.; Zalevsky, A. O.; Mészáros, B.; Bryant, P.; Good, L. L.; Laskowski, R. A.; Pozzati, G.; Shenoy, A.; Zhu, W.; Kundrotas, P.; Ruiz Serra, V.; Rodrigues, C. H. M.; Dunham, A. S.; Burke, D.; Borkakoti, N.; Velankar, S.; Frost, A.; Lindorff-Larsen, K.; Valencia, A.; Ovchinnikov, S.; Durairaj, J.; Ascher, D. B.; Thornton, J. M.; Davey, N. E.; Stein, A.; Elofsson, A.; Croll, T. I.; Beltrao, P., 2021.

(47) Baek, M.; DiMaio, F.; Anishchenko, I.; Dauparas, J.; Ovchinnikov, S.; Lee, G. R.; Wang, J.; Cong, Q.; Kinch, L. N.; Schaeffer, R. D.; Millan, C.; Park, H.; Adams, C.; Glassman, C. R.; DeGiovanni, A.; Pereira, J. H.; Rodrigues, A. V.; van Dijk, A. A.; Ebrecht, A. C.; Opperman, D. J.; Sagmeister, T.; Buhlheller, C.; Pavkov-Keller, T.; Rathinaswamy, M. K.; Dalwadi, U.; Yip, C. K.; Burke, J. E.; Garcia, K. C.; Grishin, N. V.; Adams, P. D.; Read, R. J.; Baker, D., Accurate prediction of protein structures and interactions using a three-track neural network. Science 2021, 373, 871–876.

(48) Jumper, J.; Evans, R.; Pritzel, A.; Green, T.; Figurnov, M.; Ronneberger, O.; Tunyasuvunakool, K.; Bates, R.; Zidek, A.; Potapenko, A.; Bridgland, A.; Meyer, C.; Kohl, S. A. A.; Ballard, A. J.; Cowie, A.; Romera-Paredes, B.; Nikolov, S.; Jain, R.; Adler, J.; Back, T.; Petersen, S.; Reiman, D.; Clancy, E.; Zielinski, M.; Steinegger, M.; Pacholska, M.; Berghammer, T.; Bodenstein, S.; Silver, D.; Vinyals, O.; Senior, A. W.; Kavukcuoglu, K.; Kohli, P.; Hassabis, D., Highly accurate protein structure prediction with AlphaFold. Nature 2021, 596, 583–589.

(49) Varadi, M.; Anyango, S.; Deshpande, M.; Nair, S.; Natassia, C.; Yordanova, G.; Yuan, D.; Stroe, O.; Wood, G.; Laydon, A.; Zidek, A.; Green, T.; Tunyasuvunakool, K.; Petersen, S.; Jumper, J.; Clancy, E.; Green, R.; Vora, A.; Lutfi, M.; Figurnov, M.; Cowie, A.; Hobbs, N.; Kohli, P.; Kleywegt, G.; Birney, E.; Hassabis, D.; Velankar, S., AlphaFold Protein Structure Database: massively expanding the structural coverage of protein-sequence space with high-accuracy models. Nucleic Acids Res 2022, 50, D439–D444.

(50) Tunyasuvunakool, K.; Adler, J.; Wu, Z.; Green, T.; Zielinski, M.; Zidek, A.; Bridgland, A.; Cowie, A.; Meyer, C.; Laydon, A.; Velankar, S.; Kleywegt, G. J.; Bateman, A.; Evans, R.; Pritzel, A.; Figurnov, M.; Ronneberger, O.; Bates, R.; Kohl, S. A. A.; Potapenko, A.; Ballard, A. J.; Romera-Paredes, B.; Nikolov, S.; Jain, R.; Clancy, E.; Reiman, D.; Petersen, S.; Senior, A. W.; Kavukcuoglu, K.; Birney, E.; Kohli, P.; Jumper, J.; Hassabis, D., Highly accurate protein structure prediction for the human proteome. Nature 2021, 596, 590–596.

(51) Saraiva, L. R.; Riveros-McKay, F.; Mezzavilla, M.; Abou-Moussa, E. H.; Arayata, C. J.; Makhlouf, M.; Trimmer, C.; Ibarra-Soria, X.; Khan, M.; Van Gerven, L.; Jorissen, M.; Gibbs, M.; O’Flynn, C.; McGrane, S.; Mombaerts, P.; Marioni, J. C.; Mainland, J. D.; Logan, D. W., A transcriptomic atlas of mammalian olfactory mucosae reveals an evolutionary influence on food odor detection in humans. Sci Adv 2019, 5, eaax0396.

(52) Marcinek, P.; Haag, F.; Geithe, C.; Krautwurst, D., An evolutionary conserved olfactory receptor for foodborne and semiochemical alkylpyrazines. FASEB J 2021, 35, e21638.

(53) Kurtz, R.; Steinberg, L. G.; Betcher, M.; Fowler, D.; Shepard, B. D., The Sensing Liver: Localization and Ligands for Hepatic Murine Olfactory and Taste Receptors. Front Physiol 2020, 11, 574082.

(54) Migita, K.; Iiduka, T.; Tsukamoto, K.; Sugiura, S.; Tanaka, G.; Sakamaki, G.; Yamamoto, Y.; Takeshige, Y.; Miyazawa, T.; Kojima, A.; Nakatake, T.; Okitani, A.; Matsuishi, M., Retort beef aroma that gives preferable properties to canned beef products and its aroma components. Anim Sci J 2017, 88, 2050–2056.

(55) Hou, L.; Zhang, Y.; Wang, X., Characterization of the Volatile Compounds and Taste Attributes of Sesame Pastes Processed at Different Temperatures. J Oleo Sci 2019, 68, 551–558.

(56) Henning, C.; Glomb, M. A., Pathways of the Maillard reaction under physiological conditions. Glycoconj J 2016, 33, 499–512.

(57) Bohman, B.; Phillips, R. D.; Menz, M. H.; Berntsson, B. W.; Flematti, G. R.; Barrow, R. A.; Dixon, K. W.; Peakall, R., Discovery of pyrazines as pollinator sex pheromones and orchid semiochemicals: implications for the evolution of sexual deception. New Phytol 2014, 203, 939–52.

(58) Silva-Junior, E. A.; Ruzzini, A. C.; Paludo, C. R.; Nascimento, F. S.; Currie, C. R.; Clardy, J.; Pupo, M. T., Pyrazines from bacteria and ants: convergent chemistry within an ecological niche. Sci Rep 2018, 8, 2595.

(59) Osada, K.; Kurihara, K.; Izumi, H.; Kashiwayanagi, M., Pyrazine analogues are active components of wolf urine that induce avoidance and freezing behaviours in mice. PLoS One 2013, 8, e61753.

(60) Osada, K.; Miyazono, S.; Kashiwayanagi, M., The scent of wolves: pyrazine analogs induce avoidance and vigilance behaviors in prey. Front Neurosci 2015, 9, 363.

(61) Osada, K.; Miyazono, S.; Kashiwayanagi, M., Structure-Activity Relationships of Alkylpyrazine Analogs and Fear-Associated Behaviors in Mice. J Chem Ecol 2017, 43, 263–272.

(62) Regnier, F. E., Semiochemical--structure and function. Biol Reprod 1971, 4, 309–26.

(63) Di Pizio, A.; Niv, M. Y., Computational Studies of Smell and Taste Receptors. Israel Journal of Chemistry 2014, 54, 1205–1218.

(64) Forrest, L. R.; Tang, C. L.; Honig, B., On the accuracy of homology modeling and sequence alignment methods applied to membrane proteins. Biophys J 2006, 91, 508–17.

(65) de March, C. A.; Topin, J.; Bruguera, E.; Novikov, G.; Ikegami, K.; Matsunami, H.; Golebiowski, J., Odorant Receptor 7D4 Activation Dynamics. Angew Chem Int Ed Engl 2018, 57, 4554–4558.

(66) Binder, J. L.; Berendzen, J.; Stevens, A. O.; He, Y.; Wang, J.; Dokholyan, N. V.; Oprea, T. I., AlphaFold illuminates half of the dark human proteins. Curr Opin Struct Biol 2022, 74, 102372.

(67) Ibrahim, P.; Wifling, D.; Clark, T., Universal Activation Index for Class A GPCRs. J Chem Inf Model 2019, 59, 3938–3945.

(68) Heo, L.; Feig, M., Multi-State Modeling of G-protein Coupled Receptors at Experimental Accuracy. 2022.

(69) He, X. H.; You, C. Z.; Jiang, H. L.; Jiang, Y.; Xu, H. E.; Cheng, X., AlphaFold2 versus experimental structures: evaluation on G protein-coupled receptors. Acta Pharmacol Sin 2022.

(70) Nicoli, A.; Dunkel, A.; Giorgino, T.; de Graaf, C.; Di Pizio, A., Classification Model for the Second Extracellular Loop of Class A GPCRs. J Chem Inf Model 2022, 62, 511–522.

(71) Yu, Y.; Ma, Z.; Pacalon, J.; Xu, L.; Li, W.; Belloir, C.; Topin, J.; Briand, L.; Golebiowski, J.; Cong, X., Extracellular loop 2 of G protein-coupled olfactory receptors is critical for odorant recognition. J Biol Chem 2022, 298, 102331.

(72) Woolley, M. J.; Conner, A. C., Understanding the common themes and diverse roles of the second extracellular loop (ECL2) of the GPCR super-family. Mol Cell Endocrinol 2017, 449, 3–11.

(73) Wink, L. H.; Baker, D. L.; Cole, J. A.; Parrill, A. L., A benchmark study of loop modeling methods applied to G protein-coupled receptors. J Comput Aided Mol Des 2019, 33, 573–595.

(74) Won, J.; Lee, G. R.; Park, H.; Seok, C., GalaxyGPCRloop: Template-Based and Ab Initio Structure Sampling of the Extracellular Loops of G-Protein-Coupled Receptors. J Chem Inf Model 2018, 58, 1234–1243.

(75) Di Pizio, A.; Waterloo, L. A. W.; Brox, R.; Lober, S.; Weikert, D.; Behrens, M.; Gmeiner, P.; Niv, M. Y., Rational design of agonists for bitter taste receptor TAS2R14: from modeling to bench and back. Cell Mol Life Sci 2020, 77, 531–542.

(76) Bender, B. J.; Gahbauer, S.; Luttens, A.; Lyu, J.; Webb, C. M.; Stein, R. M.; Fink, E. A.; Balius, T. E.; Carlsson, J.; Irwin, J. J.; Shoichet, B. K., A practical guide to large-scale docking. Nat Protoc 2021, 16, 4799–4832.

(77) Zhang, Y.; Vass, M.; Shi, D.; Abualrous, E.; Chambers, J.; Chopra, N.; Higgs, C.; Kasavajhala, K.; Li, H.; Nandekar, P.; Sato, H.; Miller, E.; Repasky, M.; Jerome, S., Benchmarking Refined and Unrefined AlphaFold2 Structures for Hit Discovery. ChemRxiv 2022.

(78) Lee, C.; Su, B. H.; Tseng, Y. J., Comparative studies of AlphaFold, RoseTTAFold and Modeller: a case study involving the use of G-protein-coupled receptors. Brief Bioinform 2022, 23.

(79) Moore, P. B.; Hendrickson, W. A.; Henderson, R.; Brunger, A. T., The protein-folding problem: Not yet solved. Science 2022, 375, 507.

(80) Callaway, E., What’s next for AlphaFold and the AI protein-folding revolution. Nature 2022, 604, 234–238.

(81) Akdel, M.; Pires, D. E. V.; Pardo, E. P.; Janes, J.; Zalevsky, A. O.; Meszaros, B.; Bryant, P.; Good, L. L.; Laskowski, R. A.; Pozzati, G.; Shenoy, A.; Zhu, W.; Kundrotas, P.; Serra, V. R.; Rodrigues, C. H. M.; Dunham, A. S.; Burke, D.; Borkakoti, N.; Velankar, S.; Frost, A.; Basquin, J.; Lindorff-Larsen, K.; Bateman, A.; Kajava, A. V.; Valencia, A.; Ovchinnikov, S.; Durairaj, J.; Ascher, D. B.; Thornton, J. M.; Davey, N. E.; Stein, A.; Elofsson, A.; Croll, T. I.; Beltrao, P., A structural biology community assessment of AlphaFold2 applications. Nat Struct Mol Biol 2022, 29, 1056–1067.

(82) Geithe, C.; Protze, J.; Kreuchwig, F.; Krause, G.; Krautwurst, D., Structural determinants of a conserved enantiomer-selective carvone binding pocket in the human odorant receptor OR1A1. Cell Mol Life Sci 2017, 74, 4209–4229.

(83) Yang, D.; Zhou, Q.; Labroska, V.; Qin, S.; Darbalaei, S.; Wu, Y.; Yuliantie, E.; Xie, L.; Tao, H.; Cheng, J.; Liu, Q.; Zhao, S.; Shui, W.; Jiang, Y.; Wang, M. W., G protein-coupled receptors: structure-and function-based drug discovery. Signal Transduct Target Ther 2021, 6, 7.

(84) Di Pizio, A.; Nicoli, A., In Silico Molecular Study of Tryptophan Bitterness. Molecules 2020, 25.

(85) Harini, K.; Sowdhamini, R., Computational Approaches for Decoding Select Odorant-Olfactory Receptor Interactions Using Mini-Virtual Screening. PLoS One 2015, 10, e0131077.

(86) Schneider, J.; Korshunova, K.; Musiani, F.; Alfonso-Prieto, M.; Giorgetti, A.; Carloni, P., Predicting ligand binding poses for low-resolution membrane protein models: Perspectives from multiscale simulations. Biochem Biophys Res Commun 2018, 498, 366–374.

(87) Dunkel, A.; Hofmann, T.; Di Pizio, A., In Silico Investigation of Bitter Hop-Derived Compounds and Their Cognate Bitter Taste Receptors. J Agric Food Chem 2020, 68, 10414–10423.

(88) Man, O.; Gilad, Y.; Lancet, D., Prediction of the odorant binding site of olfactory receptor proteins by human-mouse comparisons. Protein Sci 2004, 13, 240–54.

(89) Abaffy, T.; Malhotra, A.; Luetje, C. W., The molecular basis for ligand specificity in a mouse olfactory receptor: a network of functionally important residues. J Biol Chem 2007, 282, 1216–24.

(90) Ahmed, L.; Zhang, Y.; Block, E.; Buehl, M.; Corr, M. J.; Cormanich, R. A.; Gundala, S.; Matsunami, H.; O’Hagan, D.; Ozbil, M.; Pan, Y.; Sekharan, S.; Ten, N.; Wang, M.; Yang, M.; Zhang, Q.; Zhang, R.; Batista, V. S.; Zhuang, H., Molecular mechanism of activation of human musk receptors OR5AN1 and OR1A1 by (R)-muscone and diverse other musk-smelling compounds. Proc Natl Acad Sci U S A 2018, 115, E3950–E3958.

(91) Baud, O.; Etter, S.; Spreafico, M.; Bordoli, L.; Schwede, T.; Vogel, H.; Pick, H., The mouse eugenol odorant receptor: structural and functional plasticity of a broadly tuned odorant binding pocket. Biochemistry 2011, 50, 843–53.

(92) Baud, O.; Yuan, S.; Veya, L.; Filipek, S.; Vogel, H.; Pick, H., Exchanging ligand-binding specificity between a pair of mouse olfactory receptor paralogs reveals odorant recognition principles. Sci Rep 2015, 5, 14948.

(93) de March, C. A.; Yu, Y.; Ni, M. J.; Adipietro, K. A.; Matsunami, H.; Ma, M.; Golebiowski, J., Conserved Residues Control Activation of Mammalian G Protein-Coupled Odorant Receptors. Journal of the American Chemical Society 2015, 137, 8611–6.

(94) Katada, S.; Hirokawa, T.; Oka, Y.; Suwa, M.; Touhara, K., Structural basis for a broad but selective ligand spectrum of a mouse olfactory receptor: mapping the odorant-binding site. J Neurosci 2005, 25, 1806–15.

(95) Di Pizio, A.; Behrens, M.; Krautwurst, D., Beyond the Flavour: The Potential Druggability of Chemosensory G Protein-Coupled Receptors. Int J Mol Sci 2019, 20, 1402.

(96) Di Pizio, A.; Shy, N.; Behrens, M.; Meyerhof, W.; Niv, M. Y., Molecular Features Underlying Selectivity in Chicken Bitter Taste Receptors. Front Mol Biosci 2018, 5, 6.

(97) Terwilliger, T. C.; Liebschner, D.; Croll, T. I.; Williams, C. J.; McCoy, A. J.; Poon, B. K.; Afonine, P. V.; Oeffner, R. D.; Richardson, J. S.; Read, R. J.; Adams, P. D., AlphaFold predictions: great hypotheses but no match for experiment. 2022.

(98) Czerny, M.; Wagner, R.; Grosch, W., Detection of Odor-Active Ethenylalkylpyrazines in Roasted Coffee. Journal of Agricultural and Food Chemistry 1996, 44, 3268–3272.

(99) Frank, O.; Kreissl, J. K.; Daschner, A.; Hofmann, T., Accurate determination of reference materials and natural isolates by means of quantitative (1)h NMR spectroscopy. J Agric Food Chem 2014, 62, 2506–15.

(100) Porcelli, C.; Kreissl, J.; Steinhaus, M., Enantioselective synthesis of tri-deuterated (-)-geosmin to be used as internal standard in quantitation assays. J Labelled Comp Radiopharm 2020, 63, 476–481.

(101) Noe, F.; Frey, T.; Fiedler, J.; Geithe, C.; Nowak, B.; Krautwurst, D., IL-6-HaloTag((R)) enables live cell plasma membrane staining, flow cytometry, functional expression, and de-orphaning of recombinant odorant receptors. J Biol Methods 2017, 4, e81.

(102) Noe, F.; Geithe, C.; Fiedler, J.; Krautwurst, D., A bi-functional IL-6-HaloTag((R)) as a tool to measure the cell-surface expression of recombinant odorant receptors and to facilitate their activity quantification. J Biol Methods 2017, 4, e82.

(103) Graham, F. L.; Smiley, J.; Russell, W. C.; Nairn, R., Characteristics of a human cell line transformed by DNA from human adenovirus type 5. J Gen Virol 1977, 36, 59–74.

(104) Geithe, C.; Andersen, G.; Malki, A.; Krautwurst, D., A Butter Aroma Recombinate Activates Human Class-I Odorant Receptors. J Agric Food Chem 2015, 63, 9410–20.

(105) Saito, H.; Kubota, M.; Roberts, R. W.; Chi, Q.; Matsunami, H., RTP family members induce functional expression of mammalian odorant receptors. Cell 2004, 119, 679–91.

(106) Shirokova, E.; Schmiedeberg, K.; Bedner, P.; Niessen, H.; Willecke, K.; Raguse, J. D.; Meyerhof, W.; Krautwurst, D., Identification of specific ligands for orphan olfactory receptors. G protein-dependent agonism and antagonism of odorants. J Biol Chem 2005, 280, 11807–15.

(107) Jones, D. T.; Reed, R. R., Golf: an olfactory neuron specific-G protein involved in odorant signal transduction. Science 1989, 244, 790–5.

(108) Li, F.; Ponissery-Saidu, S.; Yee, K. K.; Wang, H.; Chen, M. L.; Iguchi, N.; Zhang, G.; Jiang, P.; Reisert, J.; Huang, L., Heterotrimeric G protein subunit Ggamma13 is critical to olfaction. J Neurosci 2013, 33, 7975–84.

(109) Binkowski, B.; Fan, F.; Wood, K., Engineered luciferases for molecular sensing in living cells. Curr Opin Biotechnol 2009, 20, 14–8.

(110) DeLean, A.; Munson, P. J.; Rodbard, D., Simultaneous analysis of families of sigmoidal curves: application to bioassay, radioligand assay, and physiological dose-response curves. Am J Physiol 1978, 235, e97–102.

(111) Noe, F.; Geithe, C.; Fiedler, J.; Krautwurst, D., A bi-functional IL-6-HaloTag^®^ as a tool to measure the cell-surface expression of recombinant odorant receptors and to facilitate their activity quantification. J Biol Methods 2017, 4, e82.

(112) NCBI Resource Coordinators, Database Resources of the National Center for Biotechnology Information. Nucleic Acids Research 2017, 45, D12–D17.

(113) Kumar, S.; Stecher, G.; Li, M.; Knyaz, C.; Tamura, K., MEGA X: Molecular Evolutionary Genetics Analysis across Computing Platforms. Molecular Biology and Evolution 2018, 35, 1547–1549.

(114) Thompson, J. D.; Higgins, D. G.; Gibson, T. J., CLUSTAL W: improving the sensitivity of progressive multiple sequence alignment through sequence weighting, position-specific gap penalties and weight matrix choice. Nucleic Acids Res 1994, 22, 4673–80.

(115) Saitou, N.; Nei, M., The neighbor-joining method: a new method for reconstructing phylogenetic trees. Mol Biol Evol 1987, 4, 406–25.

(116) Felsenstein, J., Confidence limits on phylogenies: An approach using the bootstrap. Evolution 1985, 39, 783–791.

(117) Zuckerkandl, E.; Pauling, L., Evolutionary divergence and convergence in proteins. 1965; p 97–166.

(118) Kooistra, A. J.; Mordalski, S.; Pandy-Szekeres, G.; Esguerra, M.; Mamyrbekov, A.; Munk, C.; Keseru, G. M.; Gloriam, D. E., GPCRdb in 2021: integrating GPCR sequence, structure and function. Nucleic Acids Res 2021, 49, D335–D343.

(119) Eswar, N.; Webb, B.; Marti-Renom, M. A.; Madhusudhan, M. S.; Eramian, D.; Shen, M. Y.; Pieper, U.; Sali, A., Comparative protein structure modeling using Modeller. Curr Protoc Bioinformatics 2006, Chapter 5, Unit-5 6.

(120) Humphrey, W.; Dalke, A.; Schulten, K., VMD: Visual molecular dynamics. Journal of Molecular Graphics 1996, 14, 33–38.

(121) Mayol, E.; Garcia-Recio, A.; Tiemann, J. K. S.; Hildebrand, P. W.; Guixa-Gonzalez, R.; Olivella, M.; Cordomi, A., HomolWat: a web server tool to incorporate ‘homologous’ water molecules into GPCR structures. Nucleic Acids Res 2020, 48, W54–W59.

(122) Rodriguez-Espigares, I.; Torrens-Fontanals, M.; Tiemann, J. K. S.; Aranda-Garcia, D.; Ramirez-Anguita, J. M.; Stepniewski, T. M.; Worp, N.; Varela-Rial, A.; Morales-Pastor, A.; Medel-Lacruz, B.; Pandy-Szekeres, G.; Mayol, E.; Giorgino, T.; Carlsson, J.; Deupi, X.; Filipek, S.; Filizola, M.; Gomez-Tamayo, J. C.; Gonzalez, A.; Gutierrez-de-Teran, H.; Jimenez-Roses, M.; Jespers, W.; Kapla, J.; Khelashvili, G.; Kolb, P.; Latek, D.; Marti-Solano, M.; Matricon, P.; Matsoukas, M. T.; Miszta, P.; Olivella, M.; Perez-Benito, L.; Provasi, D.; Rios, S.; I, R. T.; Sallander, J.; Sztyler, A.; Vasile, S.; Weinstein, H.; Zachariae, U.; Hildebrand, P. W.; De Fabritiis, G.; Sanz, F.; Gloriam, D. E.; Cordomi, A.; Guixa-Gonzalez, R.; Selent, J., GPCRmd uncovers the dynamics of the 3D-GPCRome. Nat Methods 2020, 17, 777–787.

(123) Doerr, S.; Harvey, M. J.; Noe, F.; De Fabritiis, G., HTMD: High-Throughput Molecular Dynamics for Molecular Discovery. J Chem Theory Comput 2016, 12, 1845–52.

(124) Sommer, B., Membrane Packing Problems: A short Review on computational Membrane Modeling Methods and Tools. Comput Struct Biotechnol J 2013, 5, e201302014.

(125) Lomize, M. A.; Lomize, A. L.; Pogozheva, I. D.; Mosberg, H. I., OPM: orientations of proteins in membranes database. Bioinformatics 2006, 22, 623–5.

(126) Harvey, M. J.; Giupponi, G.; Fabritiis, G. D., ACEMD: Accelerating Biomolecular Dynamics in the Microsecond Time Scale. J Chem Theory Comput 2009, 5, 1632–9.

(127) Huang, J.; MacKerell, A. D., Jr., CHARMM36 all-atom additive protein force field: validation based on comparison to NMR data. J Comput Chem 2013, 34, 2135–45.

(128) Krutler, V.; van Gunsteren, W. F.; Hnenberger, P. H., A fast SHAKE algorithm to solve distance constraint equations for small molecules in molecular dynamics simulations. Journal of Computational Chemistry 2001, 22, 501–508.

(129) Essmann, U.; Perera, L.; Berkowitz, M. L.; Darden, T.; Lee, H.; Pedersen, L. G., A smooth particle mesh Ewald method. The Journal of Chemical Physics 1995, 103, 8577–8593.

(130) Bakan, A.; Meireles, L. M.; Bahar, I., ProDy: protein dynamics inferred from theory and experiments. Bioinformatics 2011, 27, 1575–7.

(131) Pedregosa, F. a. V., G., and Gramfort, A. and Michel, V., Thirion, B. and Grisel, O., Blondel, M., Prettenhofer, P., Weiss, R. and Dubourg, V., Vanderplas, J., Passos, A., Cournapeau, D., Brucher, M., Perrot, M., Duchesnay, E., Scikit-learn: Machine Learning in Python. Journal of Machine Learning Research 2011, 12, 2825–2830.

(132) Hunter, J. D., Matplotlib: A 2D Graphics Environment. Computing in Science & Engineering 2007, 9, 90–95.

(133) Friesner, R. A.; Banks, J. L.; Murphy, R. B.; Halgren, T. A.; Klicic, J. J.; Mainz, D. T.; Repasky, M. P.; Knoll, E. H.; Shelley, M.; Perry, J. K.; Shaw, D. E.; Francis, P.; Shenkin, P. S., Glide: a new approach for rapid, accurate docking and scoring. 1. Method and assessment of docking accuracy. J Med Chem 2004, 47, 1739–49.

(134) Friesner, R. A.; Murphy, R. B.; Repasky, M. P.; Frye, L. L.; Greenwood, J. R.; Halgren, T. A.; Sanschagrin, P. C.; Mainz, D. T., Extra precision glide: docking and scoring incorporating a model of hydrophobic enclosure for protein-ligand complexes. J Med Chem 2006, 49, 6177–96.

(135) Sherman, W.; Day, T.; Jacobson, M. P.; Friesner, R. A.; Farid, R., Novel procedure for modeling ligand/receptor induced fit effects. J Med Chem 2006, 49, 534–53.

(136) consortium, P., Promoting transparency and reproducibility in enhanced molecular simulations. Nat Methods 2019, 16, 670–673.

(137) Tribello, G. A.; Bonomi, M.; Branduardi, D.; Camilloni, C.; Bussi, G., PLUMED 2: New feathers for an old bird. Computer Physics Communications 2014, 185, 604–613.

(138) Bonomi, M.; Branduardi, D.; Bussi, G.; Camilloni, C.; Provasi, D.; Raiteri, P.; Donadio, D.; Marinelli, F.; Pietrucci, F.; Broglia, R. A.; Parrinello, M., PLUMED: A portable plugin for free-energy calculations with molecular dynamics. Computer Physics Communications 2009, 180, 1961–1972.

(139) Pettersen, E. F.; Goddard, T. D.; Huang, C. C.; Meng, E. C.; Couch, G. S.; Croll, T. I.; Morris, J. H.; Ferrin, T. E., UCSF ChimeraX: Structure visualization for researchers, educators, and developers. Protein Science 2020, 30, 70–82.

