## Supplementary Information for "Modeling the Orthosteric Binding Site of the G Protein-Coupled Odorant Receptor OR5K1"

‡ These authors contributed equally.

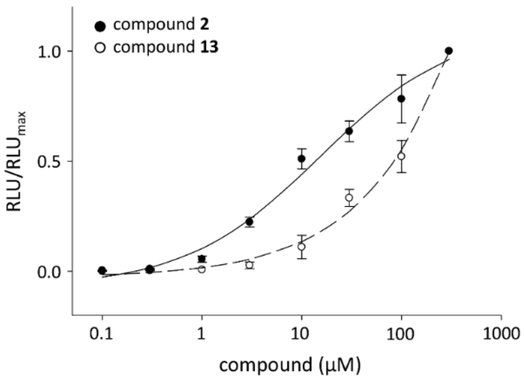


**Figure S1. Concentration-response relations of 2-ethyl-3,6-dimethylpyrazine (compound 2) and 2-ethyl-3,5-dimethylpyrazine (compound 13) on OR5K1.** Data were mock control-subtracted, normalized to the OR5K1 signal of each ligand, and displayed as mean ± SD of independent transfection experiments (n = 4). RLU = relative luminescence units.

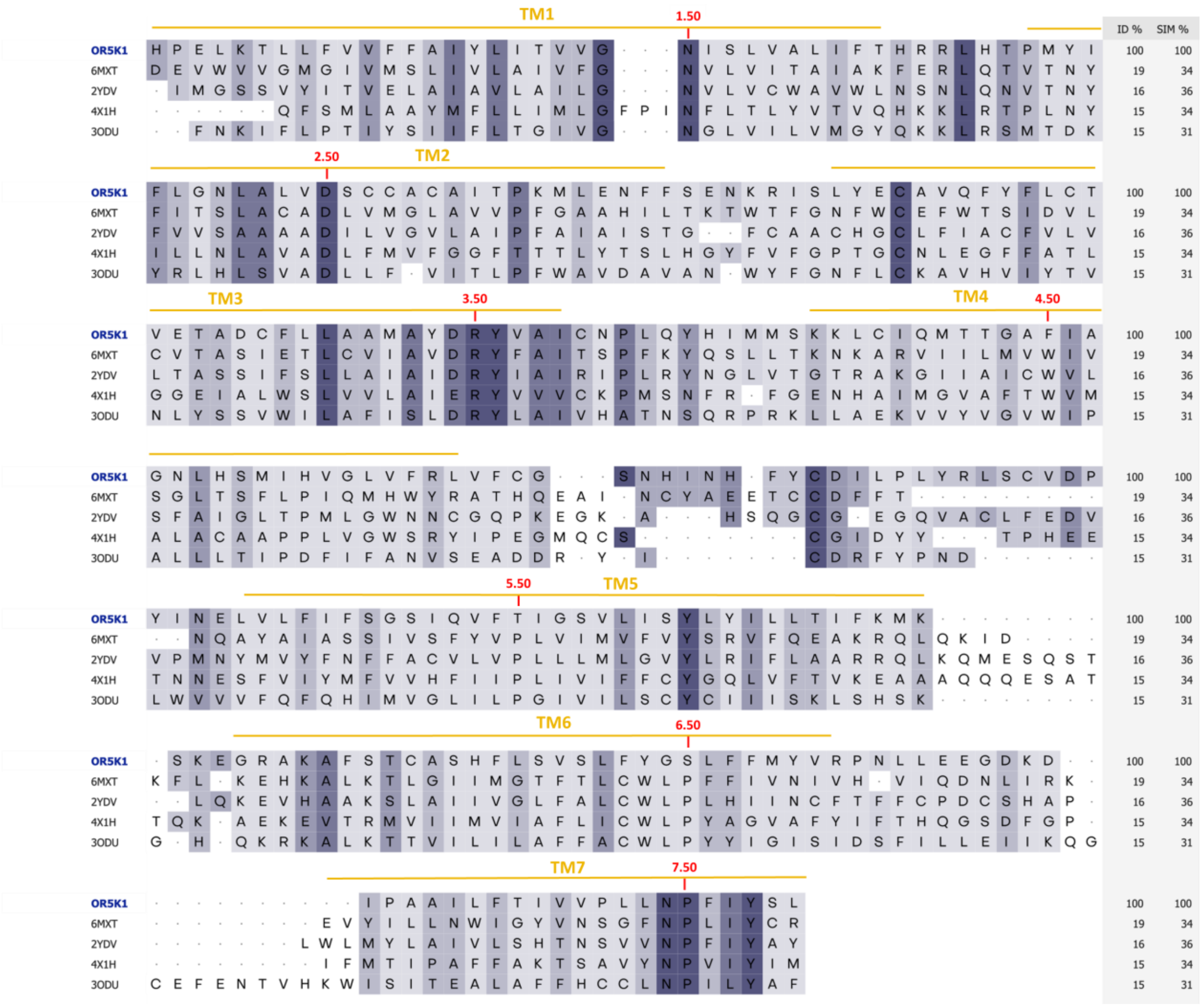

**Figure S2.** Multiple Sequence Alignment of OR5K1 (residues 20-292) to β2-adrenergic receptor (PDB ID: 6MXT), A2A receptor (PDB ID: 2YDV), Rhodopsin receptor (PDB ID: 4X1H), and CXCR4 receptor (PDB ID: 3ODU). Transmembrane (TM) domains and conserved BW positions are annotated. Template selection and sequence alignment from de March et al. 2015.(de March, Kim et al. 2015).

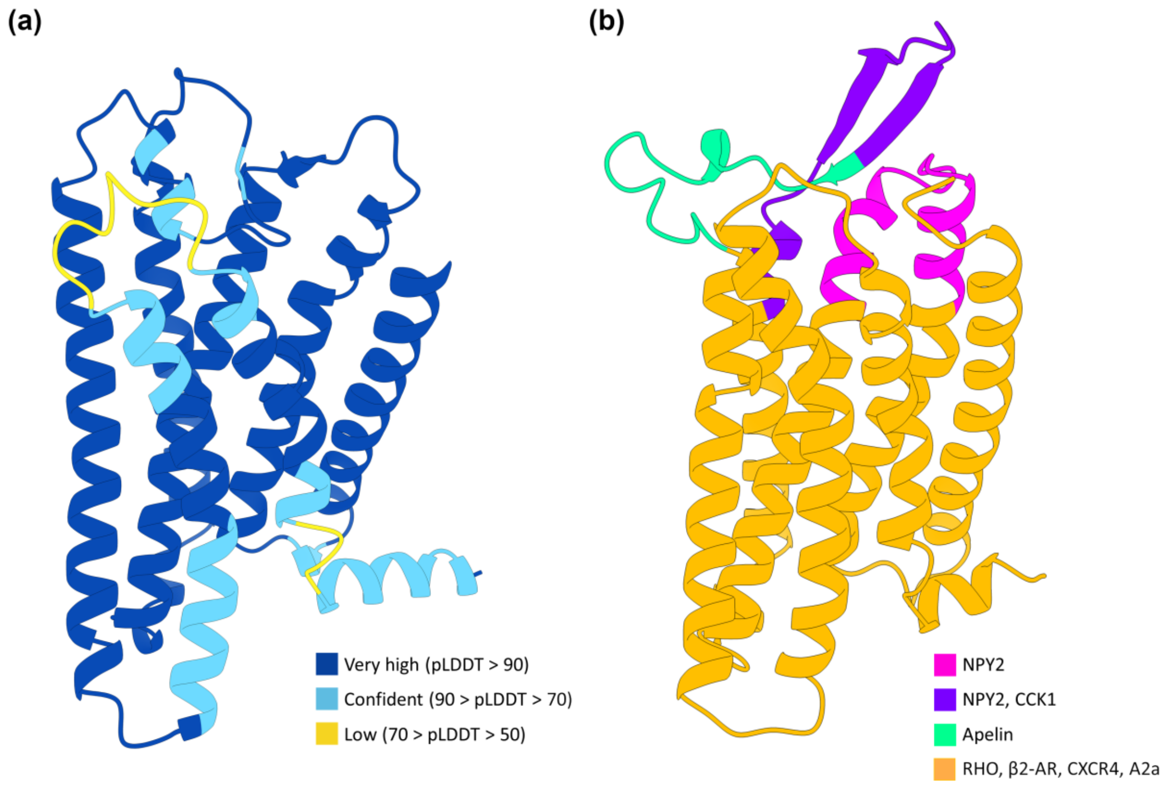


**Figure S3. OR5K1 models built with AlphaFold 2 (a) and homology modeling (b).** Alphafold 2 model (AF2) is colored by pLDDT confidence score, while OR5K1 model predicted with homology model (HM) is colored by templates coverage.

(a)

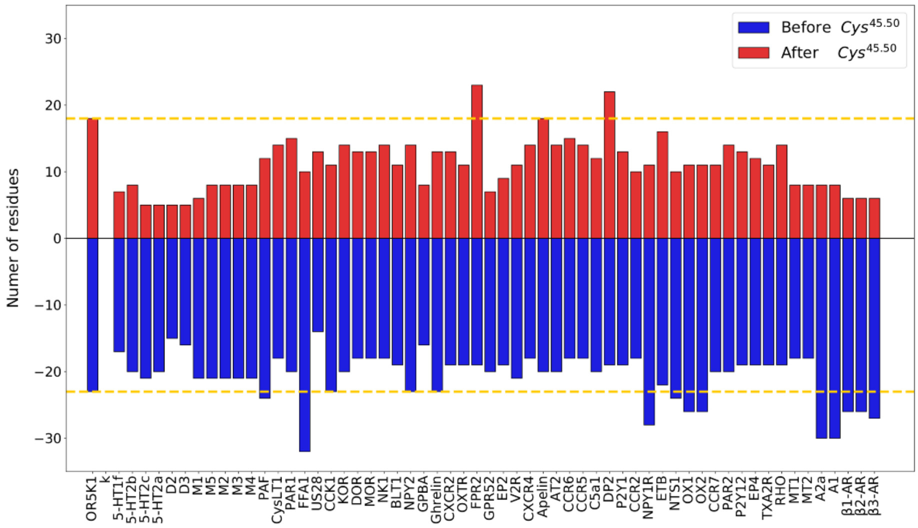


(b)

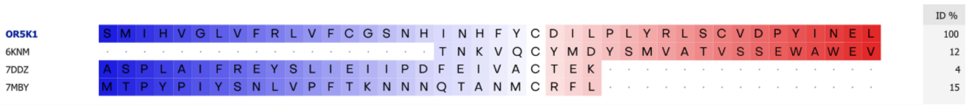


**Figure S4. (a)** Length of ECL2 for OR5K1 and experimental class A GPCRs. (**b)** Sequence Alignment of OR5K1 ECL2 region (S156$^{4.57}$- L197$^{5.37}$) to templates used for its modeling: NPY2 (PDB ID: 7DDZ) and CCK1 (PDB ID: 7MBY) before-Cys$^{45.50}$ segment and Apelin (PDB ID: 6KNM) after-Cys$^{45.50}$ segment.

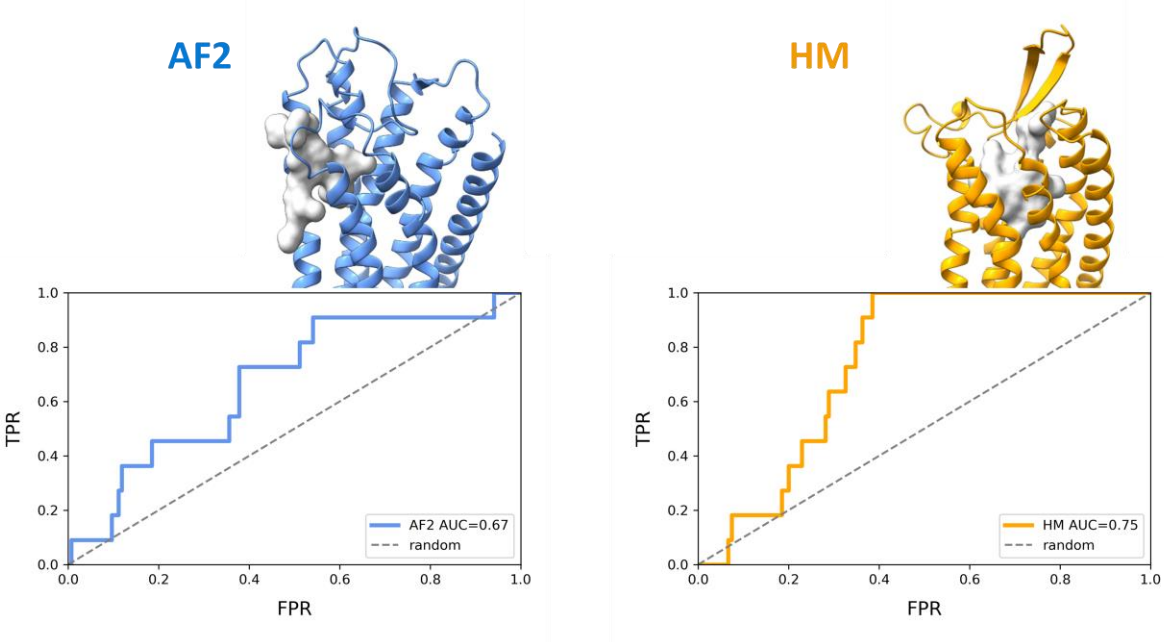


**Figure S5. Binding site and ROC analysis of the starting OR5K1 AF2 and HM models.** ROC curves plot the rate of true positives (TPR) against false positives (FPR). ROC curves are colored in blue for the AF2 model and orange for HM. Starting models are available at https://github.com/dipizio/OR5K1_binding_site.

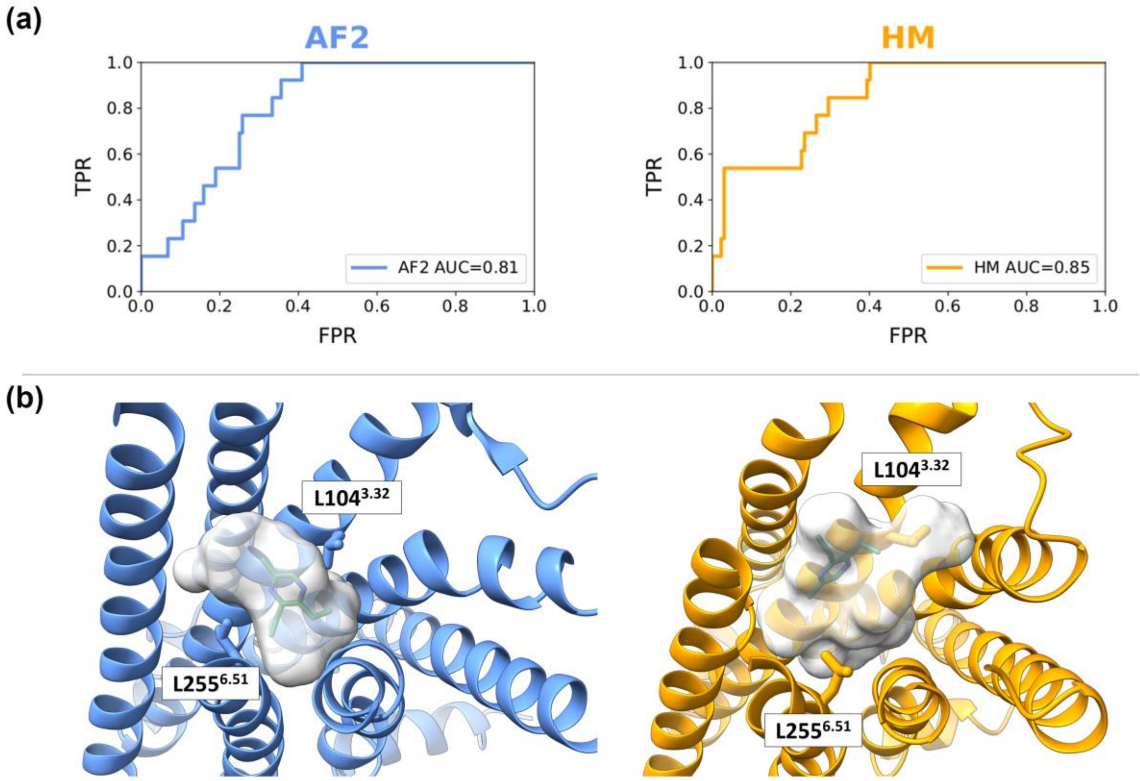


**Figure S6. (a)** ROC analysis of the OR5K1 AF2 and HM models after the first IFD simulation round. (**b**) Predicted binding modes of compound **1** within the OR5K1 binding site refined after the first IFD simulation round from AF2 and HM models.

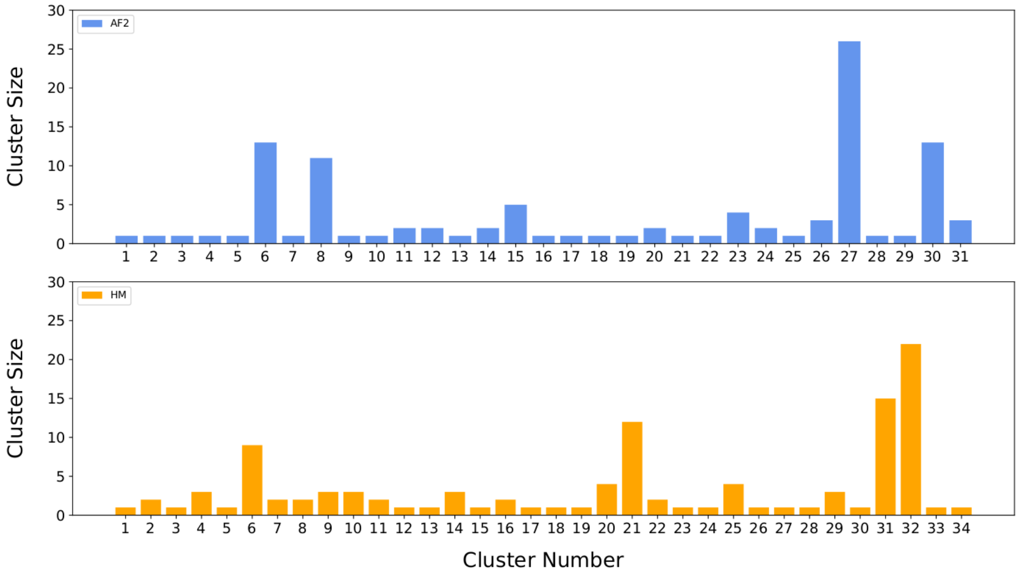


**Figure S7.** Distribution of the clusters binding poses of compound **1** in proximity to L104$^{3.32}$ and L255$^{6.51}$.

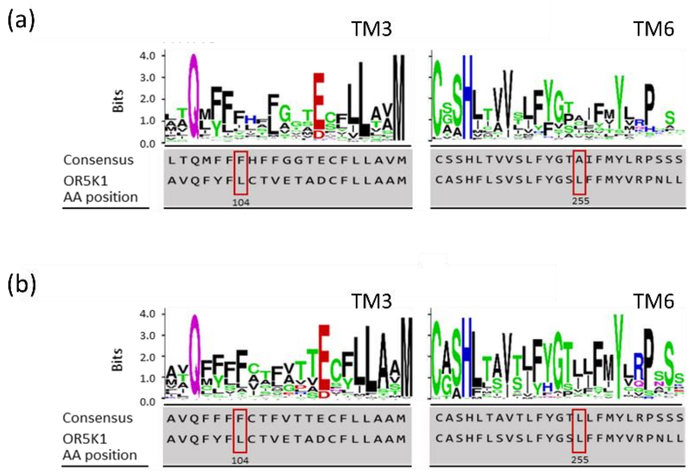


**Figure S8. Leucine residues L104$^{3.32}$ and L255$^{6.51}$ are not conserved within all human and family 5 ORs.** Alignments of TM3 and TM6 of (a) human OR5K1 and the continuing 385 human ORs as well as (b) family 5 ORs. Shown are sequence logos, the consensus sequence, and the human OR5K1 sequence with the two leucine positions (red boxes). The consensus amino acid refers to the most frequent one, which is determined by letter height and stacking order. The letters of each stack are ordered from the most frequent to the least frequent. Amino acid conservation is measured in bits, and a 100% conservation correlates to 4.32 bits (Crooks, Hon et al. 2004). Basic amino acids (K, R, H) are blue, polar (G, S, T, Y C) are green, hydrophilic (Q, N) are purple, acidic (D, E) are red, and hydrophobic (A, V, L, I, P, W, M, F) are black.

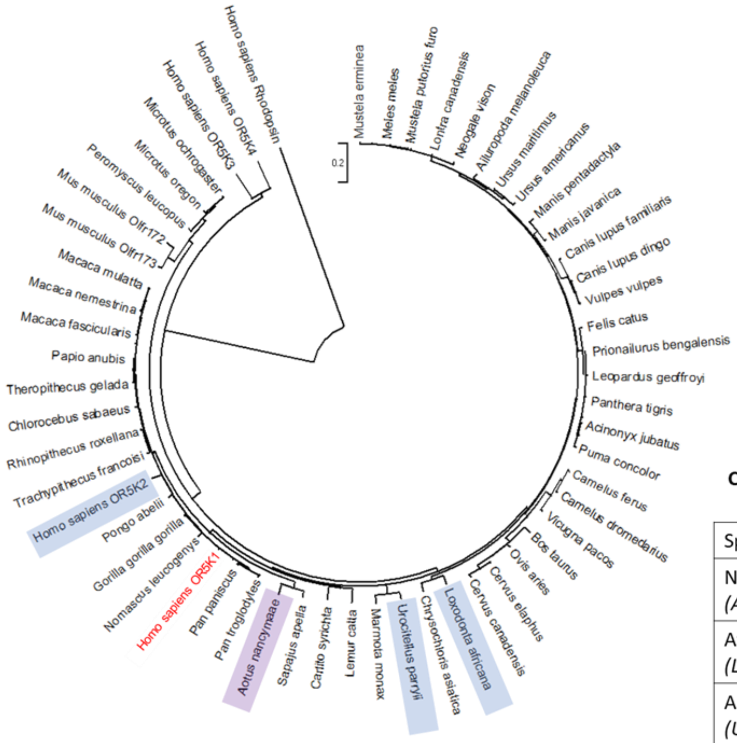


**Conservative changes at pyrazine-binding leucins in OR5K1 homologs**

| Species | L104$^{3.32}$ | L255$^{6.51}$ |
|---|---|---|
| Nancy Ma's night monkey *(Aotus nancymaae)* | Ile | Ile |
| African elephant *(Loxodonta africana)* | | Ile |
| Arctic ground squirrel *(Urocitellus parryii)* | | Ile |
| OR5K2 *(Homo sapiens)* | | Ile |

**Figure S9. Leucine residues L104$^{3.32}$ and L255$^{6.51}$ are highly conserved in OR5K1 homologs.** Circular, phylogenetic relationship of OR5K1 homologs with human rhodopsin as outgroup. Marked in purple is the species with an amino acid exchange at both conserved leucines. Highlighted in blue are the species with an amino acid exchange at L255$^{6.51}$. Homo sapiens OR5K1 is highlighted in red.

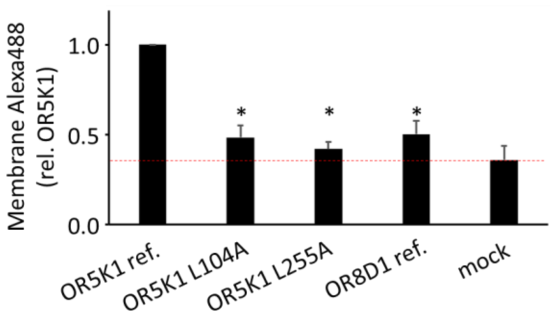


**Figure S10. Cell-surface expression of OR5K1 and the two leucine mutants.** Bar chart showing the relative surface expression of OR5K1, the two leucine mutants OR5K1 L104A and OR5K1 L255A as well as OR8D1 for comparison, using the flow cytometry assay. Red dashed line indicates the level of mock control. Data is displayed as mean ± SD (n = 3). Asterisks indicate statistically significant effects as compared to OR5K1 ref., identified by a paired t-test, with $P < 0.05$.

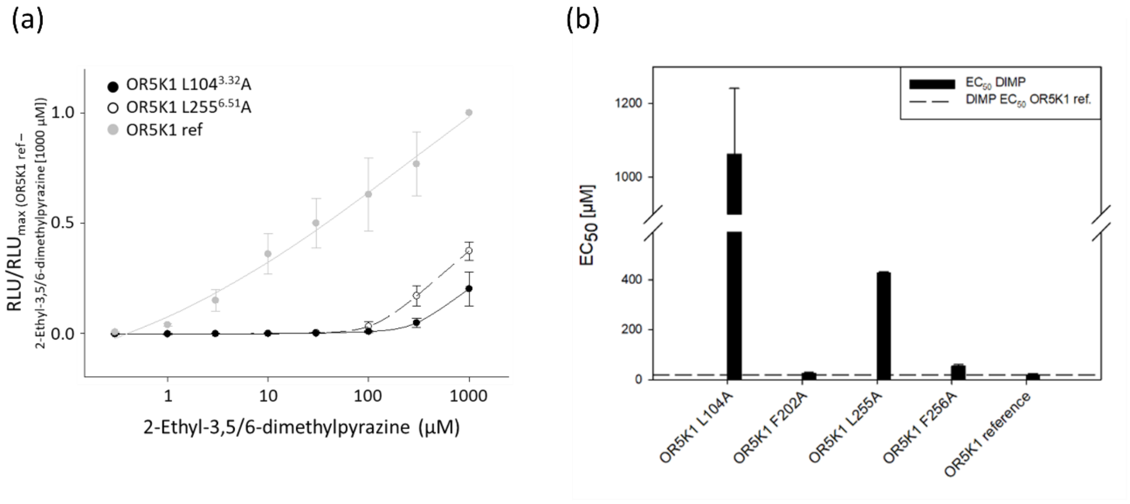


**Figure S11. (a)** Concentration–response relations of 2-ethyl-3,5/6-dimethylpyrazine on OR5K1 ref (grey circles), OR5K1 L104$^{3.32}$A (black circles), and OR5K1 L255$^{6.51}$A (white circles). Data were mock control-subtracted, normalized to the response of OR5K1 ref to 2-ethyl-3,5/6-dimethylpyrazine (1000 µM) and displayed as mean ± SD (n = 3). RLU = relative luminescence units. **(b)** Black boxes are DIMP $EC_{50}$ values for mutants L104$^{3.32}$A, L255$^{6.51}$A, F202$^{5.42}$A, and F256$^{6.52}$A and wt OR5K1; dashed line represents $EC_{50}$ of the OR5K1 reference versus 2-ethyl-3,5/6-dimethylpyrazine (DIMP). All values were derived from at least n = 3 independent experiments measured via GloSensor™ technology.

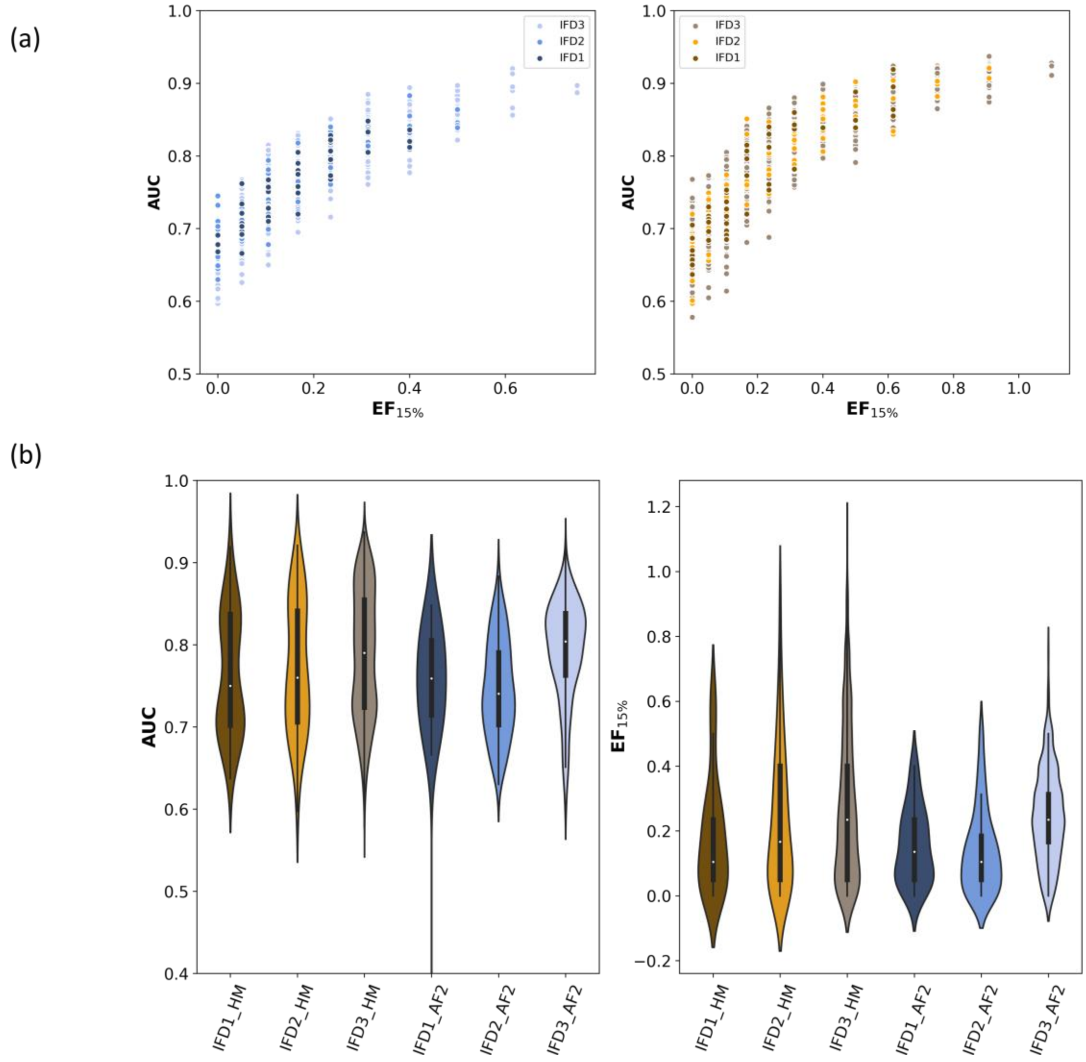


**Figure S12. (a)** Plots of EF vs. AUC values for the AF2 (blue shades) and HM models refined with IFD simulations **(b)** Distribution of AUC and EF values along the three rounds of IFD simulations.

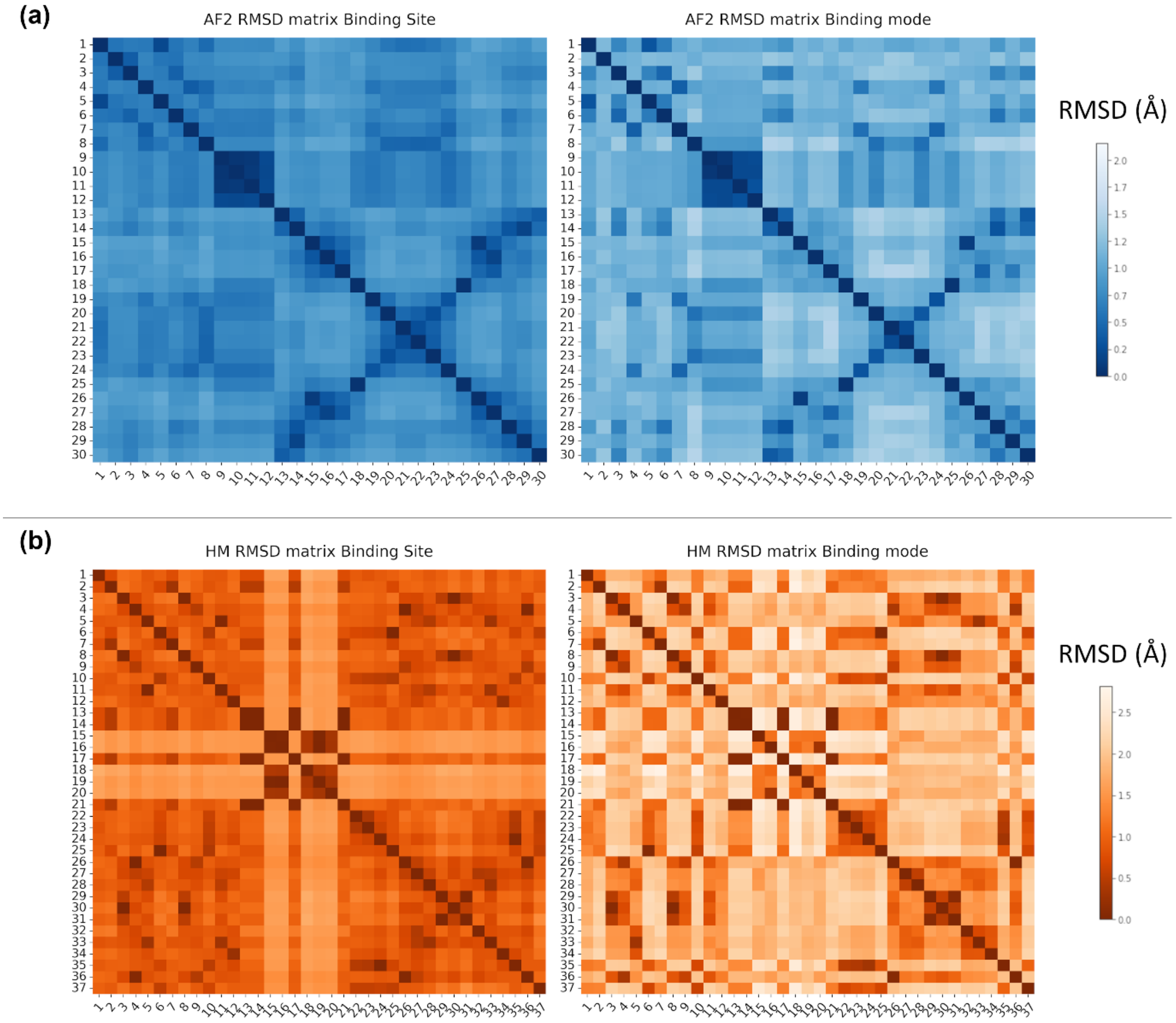


**Figure S13.** Heat maps representing the pairwise all atom RMSD matrices for the orthosteric binding site of the best performing models obtained after clustering IFD3 models. Matrices on the left refer to binding site residues, while matrices on the right include ligand coordinates in the selection of atoms for RMSD calculation. **(a)** Models from AF2. Cells are colored from dark orange to white according to increased RMSD values. Residues defining the binding site are: 104, 105, 108, 159, 199, 202, 206, 255, 256, 276, 279, 280. **(b)** Models from HM. Cells are colored from dark blue to white according to increased RMSD values. Residues defining the binding site are: 101, 104, 105, 108, 178, 180, 181, 199, 255, 258, 259, 275, 278, 279.

Models from IFD3 are available at https://github.com/dipizio/OR5K1_binding_site

**Table S1. Models from IFD1 and IFD2 with d < 0.4, AUC > 0.8**. Model names indicate if the model is resulting from the refinement of HM or AF2 models and if it is resulting from the first (IFD1) or second (IFD2) simulation round. We report AUC and $EF_{15\%}$ values, as well as information about the clustering and predicted van der Waals (vdW, kcal/mol) contribution of L104 and L255 to the binding of compound **1.** Models from IDF1 and IDF2 are available at https://github.com/dipizio/OR5K1_binding_site.

| Model | AUC | $EF_{15\%}$ | d (nm) | Cluster | Cluster Size | vdW $L104^{3.32}$ | vdW $L255^{6.51}$ |
|---|---|---|---|---|---|---|---|
| HM_IFD2_36 | 0.901 | 0.615 | 0.278 | 4 | 3 | -2.801 | -1.379 |
| HM_IFD2_1 | 0.872 | 0.400 | 0.273 | 5 | 1 | -2.477 | -0.697 |
| HM _IFD2_5 | 0.810 | 0.313 | 0.197 | 6 | 9 | -3.376 | -0.955 |
| HM_IFD2_56 | 0.834 | 0.313 | 0.281 | 8 | 2 | -2.295 | -0.768 |
| HM_IFD2_21 | 0.869 | 0.500 | 0.38 | 11 | 2 | -1.58 | -0.657 |
| HM_IFD2_20 | 0.921 | 0.909 | 0.31 | 20 | 4 | -0.329 | -2.36 |
| HM_IFD2_18 | 0.902 | 0.615 | 0.283 | 21 | 12 | -1.698 | -1.562 |
| HM_IFD2_23 | 0.864 | 0.400 | 0.391 | 23 | 1 | -1.903 | -0.385 |
| HM_IFD1_14 | 0.914 | 0.615 | 0.345 | 31 | 15 | -0.273 | -1.439 |
| HM_IFD1_3 | 0.864 | 0.615 | 0.33 | 32 | 22 | -0.221 | -1.734 |
| HM_IFD1_11 | 0.837 | 0.211 | 0.26 | 33 | 1 | -0.183 | -2.274 |
| HM_IFD1_13 | 0.855 | 0.615 | 0.399 | 34 | 1 | -0.125 | -1.503 |
| AF2_IFD2_120 | 0.833 | 0.235 | 0.285 | 8 | 11 | -2.477 | -0.697 |
| AF2_IFD2_105 | 0.883 | 0.538 | 0.356 | 15 | 5 | -1.036 | -1.246 |
| AF2_IFD2_55 | 0.808 | 0.400 | 0.265 | 21 | 1 | -2.690 | -0.937 |
| AF2_IFD2_79 | 0.855 | 0.400 | 0.153 | 27 | 26 | -2.258 | -2.669 |
| AF2_IFD2_70 | 0.864 | 0.500 | 0.149 | 30 | 13 | -2.815 | -1.221 |
| AF2_IFD2_98 | 0.841 | 0.313 | 0.216 | 31 | 3 | -2.675 | -1.284 |

**Table S2. Oligonucleotides for molecular cloning of OR5K1.**

| Gene | Oligo-nucleotide | Restriction Site | TM (°C) | | Sequence 5´→3´ |
|---|---|---|---|---|---|
| OR5K1 | al-359 | EcoRI | 58 | fw | CTGT *GAATTC* **ATG** GCT GAA GAA AAT CAT ACC ATG AAA AAT GAG TTT ATC |
| | al-360 | NotI | 59 | rv | CTGC *GCGGCCGC* **GTA** ATT TCA CAT GGA AGT TTT TGC TGC ATT TAA ATC |

TM = melting temperature, fw = forward, rv = reverse. Italic letters highlight the restriction sites. Start and Stop codons are printed in bold.

**Table S3. Vector internal oligonucleotides.**

| Vector | Oligonucleotide | TM (°C) | | Sequence 5´→3´ |
|---|---|---|---|---|
| pFN210A | 520 | 60 | fw | GTG GAC ATC GGC CCG GGT C |
| | 550 | 52 | rv | CAC AAA TAA AGC ATT TTT TTC ACT GC |

TM = melting temperature, fw = forward, rv = reverse

**Table S4. Oligonucleotides for Homo sapiens OR5K1 site directed mutagenesis.**

| Gene | Oligonucleotide | TM (°C) | | Sequence 5´→3´ |
|---|---|---|---|---|
| OR5K1 L104A | pm-275 | 57 | fw | *GTACAGTTTTATTTTGCTTGCACTGTGG* |
| | pm-276 | 57 | rv | *GTTTCCACAGTGCAAGCAAAATAAAACTG* |
| OR5K1 L255A | pm-281 | 58 | fw | *CATTATTCTATGGATCTGCTTTCTTCATGTAC* |
| | pm-282 | 58 | rv | *GTACATGAAGAAAGCAGATCCATAGAATAATG* |

TM = melting temperature, fw = forward, rv = reverse

**Table S5. NCBI reference sequences of olfactory receptor genes investigated.**

| Gene Description | Species | Common Species Name | NCBI Reference Sequence (Accession-number) |
|---|---|---|---|
| OR5K1 | *Acinonyx jubatus* | Cheetah | XP_014936391.1 |
| OR5K1 | *Ailuropoda melanoleuca* | Giant panda | XP_011231001.2 |
| OR5K1 | *Aotus nancymaae* | Nancy Ma's night monkey | XP_012332612.1 |
| OR5K1 | *Bos taurus* | Cattle | NP_001377368.1 |
| OR5K1 | *Camelus dromedarius* | Dromedary | XP_010976859.1 |
| OR5K1 | *Camelus ferus* | Wild Bactrian camel | XP_006181138.2 |
| OR5K1 | *Canis lupus dingo* | Australian dingo | XP_025272970.1 |
| OR5K1 | *Canis lupus familiaris* | Domestic dog | NP_001376061.1 |
| OR5K1 | *Carlito syrichta* | Philippine tarsier | XP_008064628.1 |
| OR5K1 | *Cervus canadensis* | Elk | XP_043304847.1 |
| OR5K1 | *Cervus elaphus* | Red deer | XP_043748867.1 |

| | | | |
|---|---|---|---|
| OR5K1 | *Chlorocebus sabaeus* | Green monkey | XP_007984292.2 |
| OR5K1-like | *Chrysochloris asiatica* | Cape golden mole | XP_006868660.1 |
| OR5K1 | *Felis catus* | Domestic cat | XP_003991608.3 |
| OR5K1 | *Gorilla gorilla gorilla* | Western lowland gorilla | XP_004036007.1 |
| OR5K1 | *Homo sapiens* | Human | NP_001004736.2 |
| OR5K2 | *Homo sapiens* | Human | NP_001004737.1 |
| OR5K3 | *Homo sapiens* | Human | NP_001005516.1 |
| OR5K4 | *Homo sapiens* | Human | NP_001005517.1 |
| OR5K1 | *Lemur catta* | Ring-tailed lemur | XP_045396378.1 |
| OR5K1 | *Leopardus geoffroyi* | Geoffroy's cat | XP_045293055.1 |
| OR5K1 | *Lontra canadensis* | North American river otter | XP_032708517.1 |
| OR5K1 | *Loxodonta africana* | African bush elephant | XP_003418985.1 |
| OR5K1 | *Macaca fascicularis* | Crab-eating macaque | XP_045241289.1 |
| OR5K1 | *Macaca mulatta* | Rhesus macaque | NP_001180719.3 |
| OR5K1 | *Macaca nemestrina* | Southern pig-tailed macaque | XP_011732248.1 |
| OR5K1 | *Manis javanica* | Sunda pangolin | XP_036864239.1 |
| OR5K1 | *Manis pentadactyla* | Chinese pangolin | XP_036772454.1 |
| OR5K1 | *Marmota monax* | Groundhog | XP_046311290.1 |
| OR5K1 | *Meles meles* | European badger | XP_045859641.1 |
| OR5K1 | *Microtus ochrogaster* | Prairie vole | XP_005344986.1 |
| OR5K1 | *Microtus oregon* | Stoat | XP_041528659.1 |
| Olfr172 | *Mus musculus* | Mouse | NP_667212.2 |
| Olfr173 | *Mus musculus* | Mouse | NP_667211.2 |
| OR5K1 | *Mustela erminea* | Ermine | XP_032200493.1 |
| OR5K1 | *Mustela putoriusfuro* | Ferret | XP_004772048.1 |
| OR5K1 | *Neogale vison* | American mink | XP_044110727.1 |
| OR5K1 | *Nomascus leucogenys* | Northern white-cheeked gibbon | XP_003261757.2 |
| OR5K1 | *Ovis aries* | Domestic sheep | XP_004002914.1 |
| OR5K1 | *Pan paniscus* | Bonobo | XP_003821943.1 |
| OR5K1 | *Panthera tigris* | Tiger | XP_007096003.2 |
| OR5K1 | *Pan troglodytes* | Common chimpanzee | XP_526253.2 |
| OR5K1 | *Papio anubis* | Olive baboon | XP_031518815.1 |
| OR5K1 | *Peromyscus leucopus* | White-footed mouse | XP_028723391.1 |
| OR5K1 | *Pongo abelii* | Sumatran orangutan | XP_002813420.2 |
| OR5K1 | *Prionailurus bengalensis* | Leopard cat | XP_043451993.1 |
| OR5K1 | *Puma concolor* | Cougar | XP_025769507.1 |
| OR5K1 | *Rhinopithecus roxellana* | Golden snub-nosed monkey | XP_010352038.2 |
| OR5K1 | *Sapajus apella* | Tufted capuchin | XP_032154801.1 |
| OR5K1 | *Theropithecus gelada* | Gelada | XP_025233483.1 |
| OR5K1 | *Trachypithecus francoisi* | François' langur | XP_033067425.1 |
| OR5K1 | *Urocitellus parryii* | Arctic ground squirrel | XP_026258216.1 |
| OR5K1 | *Ursus americanus* | American black bear | XP_045641565.1 |
| OR5K1 | *Ursus maritimus* | Polar bear | XP_008700260.1 |
| OR5K1 | *Vicugna pacos* | Alpaca | XP_006208125.1 |
| OR5K1 | *Vulpes vulpes* | Redfox | XP_025846410.1 |

**References:**

Crooks, G. E., G. Hon, J.-M. Chandonia and S. E. Brenner (2004). "WebLogo: a sequence logo generator." Genome Research **14**(6): 1188-1190.

de March, C. A., S. K. Kim, S. Antonczak, W. A. Goddard, 3rd and J. Golebiowski (2015). "G protein-coupled odorant receptors: From sequence to structure." Protein Sci **24**(9): 1543-1548.
